# Community-level physiological profiling of carbon substrate metabolization by microbial communities associated to sediments and water in karstic caves from Romania

**DOI:** 10.1101/2025.02.17.638648

**Authors:** Diana Felicia Panait, Andrei Marian Panait, Adorján Cristea, Erika Andrea Levei, Oana Teodora Moldovan, Horia Leonard Banciu

**Affiliations:** Institute for Research, Development in Applied Natural Sciences, Babeș-Bolyai University, Cluj-Napoca, Romania; Electron Microscopy Center, Faculty of Biology and Geology, Babeș-Bolyai University, Cluj-Napoca, Romania; Doctoral School of Integrative Biology, Faculty of Biology and Geology, Babeș-Bolyai University, Cluj-Napoca, Romania; S.C. Brillio Romania SRL, Cluj-Napoca, Romania Adorján Cristea (AC); Department of Ecology and Taxonomy, Faculty of Biology and Geology, Babeș-Bolyai University, Cluj-Napoca, Romania; INCDO-INOE 2000, Research Institute for Analytical Instrumentation, Cluj-Napoca, Romania; Emil Racovita Institute of Speleology, Cluj-Napoca Department, Cluj-Napoca, Romania; Romanian Institute of Science and Technology, Cluj-Napoca, Romania; Centro Nacional de Investigación sobre la Evolución Humana, CENIEH, Burgos, Spain; Department of Molecular Biology and Biotechnology, Faculty of Biology and Geology, Babeș-Bolyai University

**Author notes:** Corresponding author: Diana Felicia Panait (DFP), Horia Leonard Banciu (HLB).

**Keywords:** caves, Community-Level Physiological Profile, Generalized Additive Models, metabolic diversity, organic carbon substrates, sediments, water, moonmilk

## Abstract

Cave ecosystems comprise specialized microbial communities that play essential roles in biogeochemical cycles; yet their metabolic capabilities and ecological functions are not yet fully understood. As conventional cultivation techniques offer limited insights into the metabolic capabilities, methods based on direct functionality screening could provide more in-depth knowledge of cave microbial activity. In this study, we utilized the Community-Level Physiological Profiling (CLPP) based on Biolog® EcoPlate™ approach to assessing carbon substrate utilization by microbial communities associated with pool water, sediment, sediment (limon), and moonmilk from five caves in Romania. Principal Component Analysis (PCA) and Generalized Additive Models (GAMs) statistics were employed to infer the patterns of C-substrate metabolization and their environmental drivers. Environmental variables such as sodium (Na) and electrical conductivity (EC) significantly impacted C-utilization capabilities as indicated by both PCA and GAM. The latter analysis elucidated non-linear relationships between variables, such as EC, Na, and Mg, and microbial metabolic diversity indices. However, distinct C utilization patterns were detected among sampled sites and chemical types. Unlike moonmilk samples whose associated microbial communities appeared as exhibiting low C-substrate utilization, the highest activity was shown in cave pool water samples with the associated microbial communities extensively consuming D-galacturonic acid and Tween 80. Conversely, substrates like L-threonine and α-ketobutyric acid showed limited utilization across all cave samples. Average Well Color Development (AWCD) and Shannon diversity indices indicated that microbial communities associated to samples from Cloșani and Muierilor caves demonstrated the highest metabolic diversity. Our findings suggested that metabolic profiling using Biolog®EcoPlates™ method combined with multivariate statistical methods might prove as suitable approach to effectively screen for cave microbial functionality and the probable environmental drivers. Besides, this work distinguishes from similar studies by relying on GAM analysis to predict the environmental factors governing the microbially-mediated organic carbon degradation in subterranean ecosystems.

## Introduction

The nutrient-limited subterranean ecosystems are inhabited by microorganisms with slow metabolism and growth rates (Epure et al., 2014). These cave-dwelling microbes are assumedly involved in biogeochemical cycles of major elements such as C, N, P or S. Therefore investigation of taxonomic and metabolic diversity is crucial in enlarging our understanding of the interactions and ecological roles of cave microorganisms (Barton and Northup., 2007). Yet, investigating the microbial diversity in cave ecosystems poses significant challenges, including the limitations of the traditional culturing techniques, which recover only a small fraction of the members of cave-inhabiting microbial communities (Jones et al., 2015). Additionally, culture-dependent techniques provide limited understanding of microbial activity *in situ*, failing to capture the complexity of functional relationships within microbial communities. This constraint interferes with accurately understanding these microorganisms’ ecological functions and dynamics in their natural environment. In this context, complementary methods aiming at unveiling the metabolic potential of microorganisms, such as Community-Level Physiological Profiles (CLPP) utilizing a rapid tool such as organic carbon substrate-based EcoPlate™ plates developed by Biolog® (Hayward, CA, USA) are essential for obtaining a broader perspective of the metabolic capabilities of cave microbial communities. This approach allows for an indirect characterization of community functionality by evaluating carbon substrate utilization patterns providing a clearer image of microorganisms’ functional diversity and ecological potential in cave environments. The Biolog® EcoPlate™ is a time– and cost-effective method to monitor the spatio-temporal carbon substrate preferences of microbial communities inhabiting a diverse range of habitats including groundwater (Melita et al., 2023), surface freshwater (Boteva et al., 2024), brackish-to-saline waters (Cristea et al., 2014), wetlands (Teng et al., 2020), agricultural soils (Rutgers et al., 2015), rhizosphere (Cacchio and Del Gallo, 2019), volcanic soils and mud (Amaresan et al., 2018; Asif et al., 2024), and limestone monuments (Andrei et al., 2017). Moreover, this versatile approach has been applied to assess the carbon utilization by bacterial communities in karst caves (Yun et al., 2018) and lava tubes ice caves (O’Connor et al., 2021). Overall, the noteworthy benefit of this method is that it is comparable to high-resolution, yet more resource-consuming molecular techniques such as metagenomics. Although the Biolog® EcoPlate™ method provides a fairly limited information on microbial metabolic capabilities, its rapidity in profiling the community-wide metabolic activity makes it a valuable complementary tool for metabolic fingerprinting thus aiding our understanding of functional diversity within cave ecosystems.

Despite increasing interest in exploring cave microbial communities, considerable gaps still exist understanding their individual and joint metabolic processes. Present limitations result from focusing on taxonomy rather than functionality, scarcity of *in situ* studies, metabolite profiling, and a poor understanding of biogeochemical processes in cave or other subterranean environments. Here, we aimed to employ the Biolog® EcoPlate™ technique to survey the carbon substrate utilization patterns of microbial communities inhabiting cave sediments, pool water and substrate (limon), and moonmilk (white precipitate with aggregates of fine carbonate crystals), from five Romanian caves, thereby addressing the diversity of carbon metabolization traits and the delineation of their main environmental drivers.

## Materials and Methods

### Sampling sites

Five karstic caves (Cloșani, Ferice, Leșu, Muierilor, and Topolnița) were sampled in this study (Figure 1). A brief description of sampled caves is provided below.

**Figure 1.**
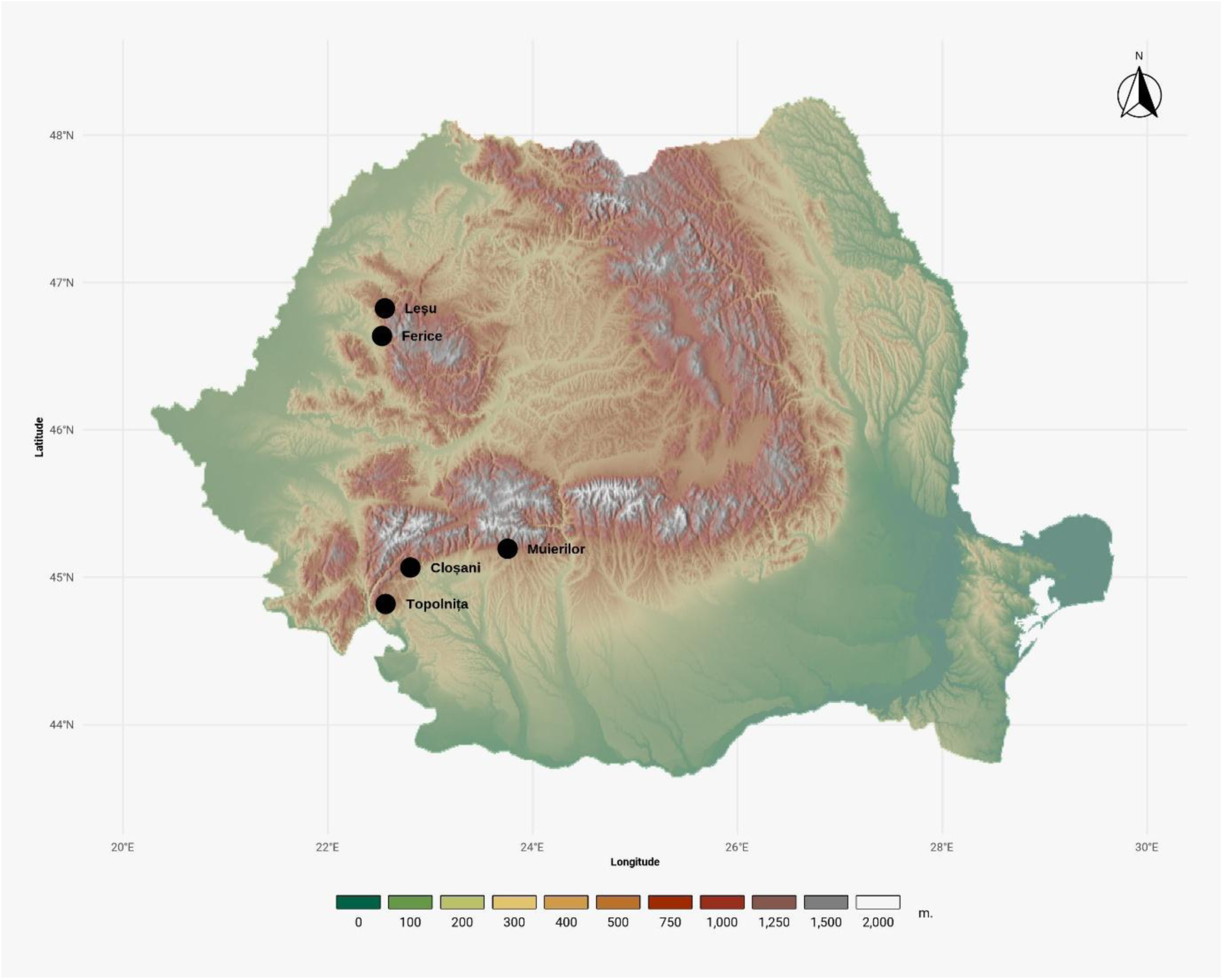
Geographic and altitudinal distribution of the Romanian sampled caves. Marked sites include Cloșani and Muierilor caves in Gorj County (Southern Romania), Topolnița Cave in Mehedinți County (SW Romania), Ferice, and Leșu caves in Bihor County (NW Romania). The map was generated using R Studio with topographic data using the ‘geodata’ package with the data source from Shuttle Radar Topography Mission (SRTM), specifically the hole-filled CGIAR-SRTM (90 m resolution) from https://srtm.csi.cgiar.org/.

#### a. Cloșani Cave

Located in Southern Carpathians, in Cloșani village, the cave has a narrow entrance found at an altitude of 443m above sea level (a.s.l). The two galleries have a length of 1100m, and temperatures of about 11°C in the depth of the cave, increased humidity and a lack of air currents (Bleahu et al., 1976).

#### b. Ferice Cave

Ferice Cave is located in North-Western Romania, Apuseni Mountains at an elevation of 410 m a.s.l. in a region with oceanic climate influences. A low-flow stream crosses the main horizontal passage, which measures around 260 meters in length. During the winter season a small number of bats frequent the cave. The temperature within the cave ranges from 11.4°C to 12.7°C, but the humidity consistently is ranging from 95% in November (its highest point) to around 80% in May with its lowest point (Moldovan et al., 2023).

#### c. Leșu Cave

The Leșu Cave (Peștera cu Apă din Valea Leșului) situated in the Apuseni Mountains is a designated protected area classified as a natural reserve (type IV IUCN). The cave system features a primary gallery approximately 1 km long, traversed by a meandering water stream for the initial 300 m, forming alluvial terraces. The annual air temperature within the cave averages between 8.5° and 10°C. The cave has a significant hibernation colony of multiple bat species, primarily near the entrance (Zoltan and Szántó, 2003; Bücs et al., 2012).

#### d. Muierilor Cave

Muierilor Cave is located in the Southern Carpathians near Baia de Fier, Gorj County and has multiple chambers within its karstic system. Muierilor Cave is located at an altitude of approximately 645 m a.s.l. and has a total length of over 8000 m. The cave system includes pristine sections and a show cave (Level 2 – accessible for visitors). It is a protected natural monument with designated scientific reserves (Mirea et al., 2021).

#### e. Topolnița Cave

The Topolnița Cave is an extensive karst system that covers an area of more than 20 km. Situated at an elevation of ∼400 m a.s.l., the cave has significant bat colonies and important guano deposits (Moldovan et al., 2023; Cleary et al., 2019).

### Sample collection and Biolog® EcoPlate™ assay

Environmental samples were collected from the studied cave sites, including cave water and substrate (limon), sediment, and moonmilk (see Table 1). Pool substrate samples were considered and processed as sediment samples. All samples were collected from areas free from human impact, including samples from show caves like Muierilor. Five grams of fresh sediment (including limon and moonmilk samples) were suspended in 50 mL NaCl 0.85% solution, and stirred for about 30 minutes on a water-bath shaker (150-200 rpm) at 23°C. After one hour of on-the-table sedimentation, 100 µL of the resulted cellular suspension was inoculated in each well from the EcoPlate™ plates (Biolog®, Hayward, CA, USA) under a laminar airflow hood, ensuring the prevention of contamination throughout all procedures. The 96-well Biolog® EcoPlate™ contains a total of 31 carbon substrates in triplicate and three wells without substrate serving as a control. One plate was used for each sample. Both the initial, and further OD measurements (at 590 nm) were assessed on a plate reader (FLUOstar® Omega. BMG Labtech, Offenburg, Germany). Plates were incubated up to 8 days (200 hours) at 16°C and specific intervals of time between readings were established based on the literature review which mostly depends on the color development on plates (Garland, 1997; Ițcuș, 2016). A 12-hour interval between measurements was established and carried out until no changes in both color development and OD_590_ values were detected.

**Table 1.**
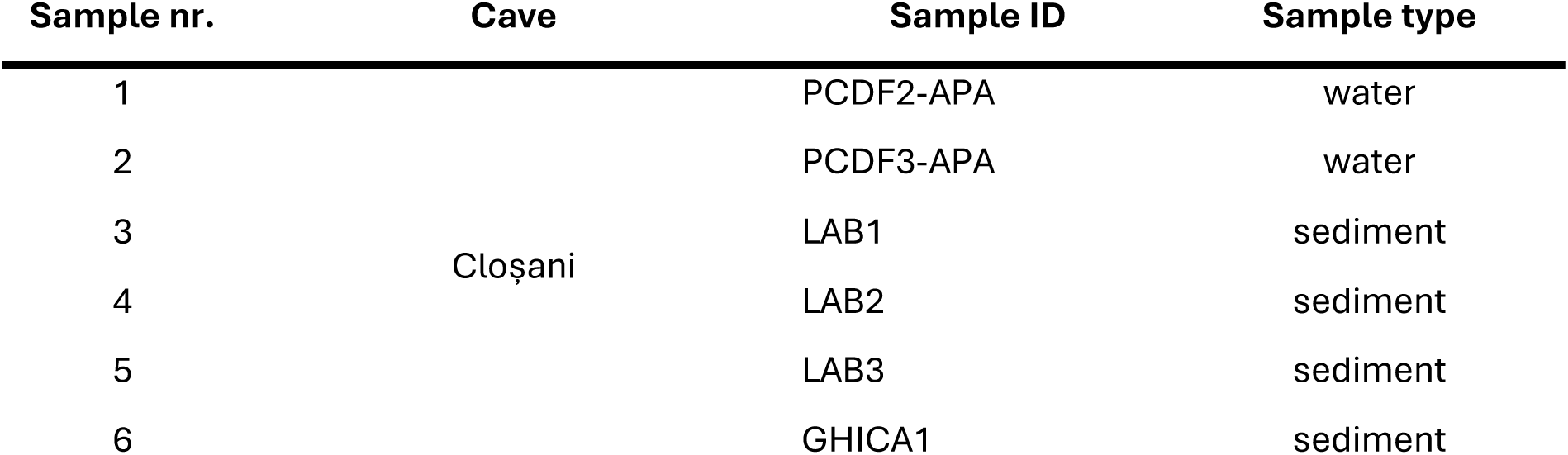
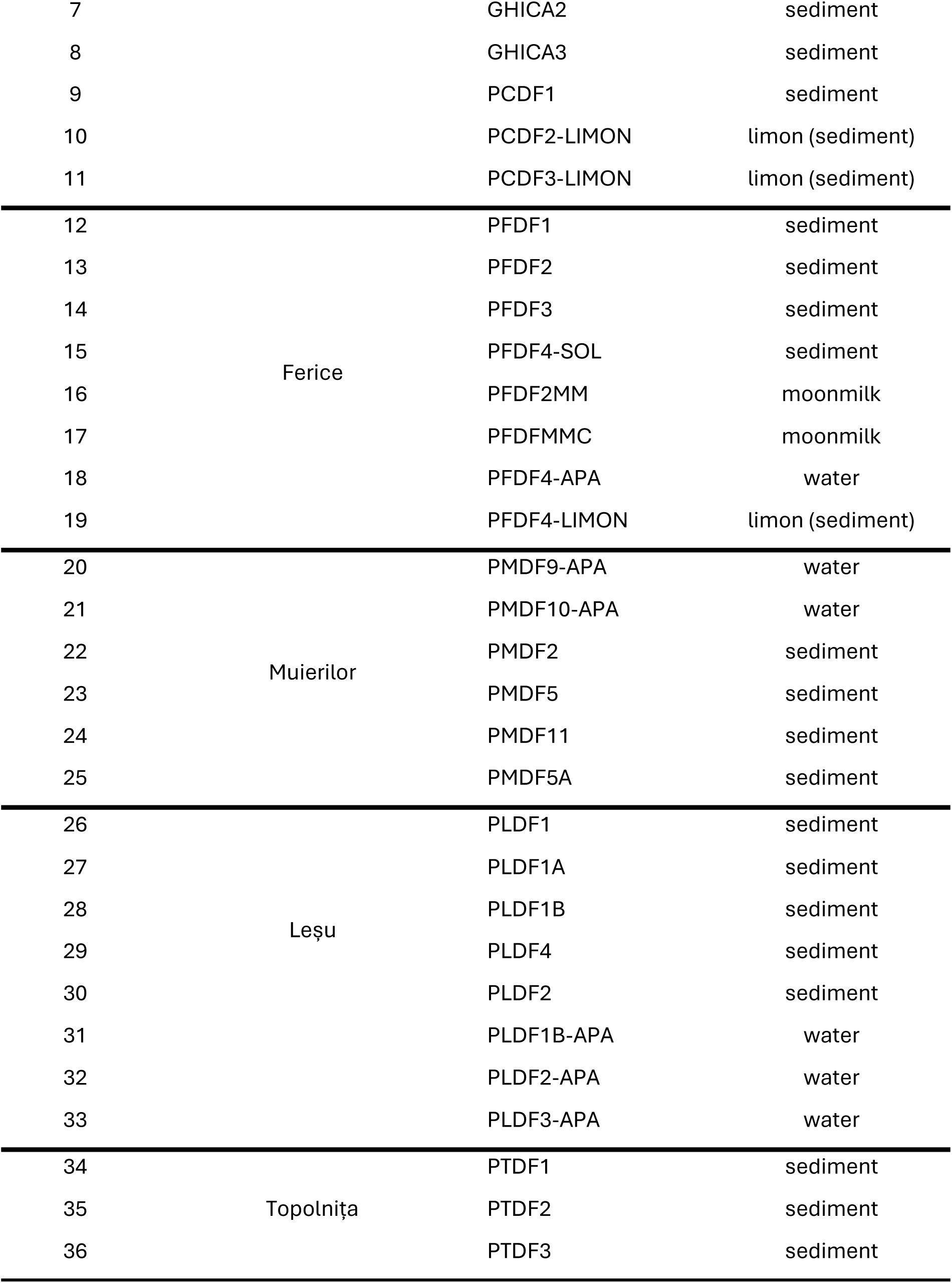
Summary of the caves and sample types used in the analysis of community-level physiological profiling.

### Physico-chemical analyses

The pH and electrical conductivity (EC) were assessed in water samples and 1/5 sediment-to-water extracts using the Seven Excellence multiparameter (Mettler Toledo, Greifensee, Switzerland). To measure the concentration of elements, one gram of dried sediment was digested with *aqua regia* (21 mL of 12 M HCl and 7 mL of 15.8 M HNO_3_), subsequently filtered and diluted to 100 mL with 0.5 M HNO_3_. The *aqua regia*-extractable fraction includes both physiologically available and unavailable metals. Water samples were acidified with 15.8 M HNO_3_ and filtered using cellulose acetate membrane filters with a pore size of 0.45 μm for the determination of dissolved metals and phosphorus contents. The concentrations of Na, Mg, K, Ca, and P in water, as well as Na, Mg, K, Ca, P, Al, Fe, S, and Mn in sediments, limon and moomilk were quantified using inductively coupled plasma optical emission spectrometry with an Optima 5300DV (Perkin Elmer, Waltham, MA, USA) spectrometer. Additionally, the concentrations of Al, Fe, As, Cr, Mn, Co, Ni, Cu, and Zn in water, along with As, Cr, Co, Ni, Cu, and Zn in sediments, were determined via inductively coupled mass spectrometry utilizing an Elan DRC II (Perkin Elmer, Waltham, MA, USA). The carbon, and nitrogen contents in sediments were quantified using a Flash 2000 CHNS/O analyzer (ThermoFisher Scientific, Waltham, MA, USA). Total nitrogen (TN) in water was quantified using catalytic combustion, followed by the oxidation of nitrogen monoxide to nitrogen dioxide using ozone, and subsequent detection via chemiluminescence with a Multi N/C 2100S Analyser (Analytik Jena, Jena, Germany). Dissolved carbon (DC) and dissolved inorganic carbon (DIC) were measured in water samples filtered via 0.45 μm PTFE syringe filters using catalytic combustion and infrared detection of CO_2_ utilizing a Multi N/C 2100S Analyser (Analytik Jena, Jena, Germany). Dissolved organic carbon (DOC) was derived by subtracting dissolved inorganic carbon (DIC) from total carbon (DC). Sulfate (SO_4_^2−^), nitrate (NO_3_^−^), chloride (Cl^−^), and phosphate (PO_4_^3−^) were quantified using ion chromatography on a 761 Compact IC (Metrohm, Switzerland).

### Data acquisition, preprocessing and processing

Data from OD_590_ measurements were extracted from individual .xlsx files using *readxl* package. Data from some samples from Lesu Cave described in Bogdan et al. (2023) were reanalyzed in a comparative context and with a more comprehensive statistical analysis. Each sample was identified by its unique sampling site, sample type and the hour of incubation. The absorbance (OD_590_) values were corrected against water blanks using designated wells and averaged across triplicate measurements for each carbon substrate. Further, the dataset was processed to remove negative values after correction.

Some samples were read up to 360 hours, but given that no changes were detected in the color development or OD_590_ measurements we excluded the timepoints after 204 hours. All analyses were performed using R (R Core Team, 2020) and key packages included the following:

- **readxl**, **dplyr** and **tidyr** for data processing;
- **mgcy** for GAMs;
- **factoextra** for PCA;
- **ggplot2** and **pheatmap** for data visualization.

### Calculation of microbial metabolic diversity metrics

Microbial metabolic diversity metrics were calculated for each sample based on the formulas and indices from the literature (Feigl et al., 2017; Zak et al., 1994) and are summarized in Table 2. The principal microbial metabolic diversity metrics provide insights into both the metabolic capacity and functional evenness of the microbial communities.

**Table 2.**
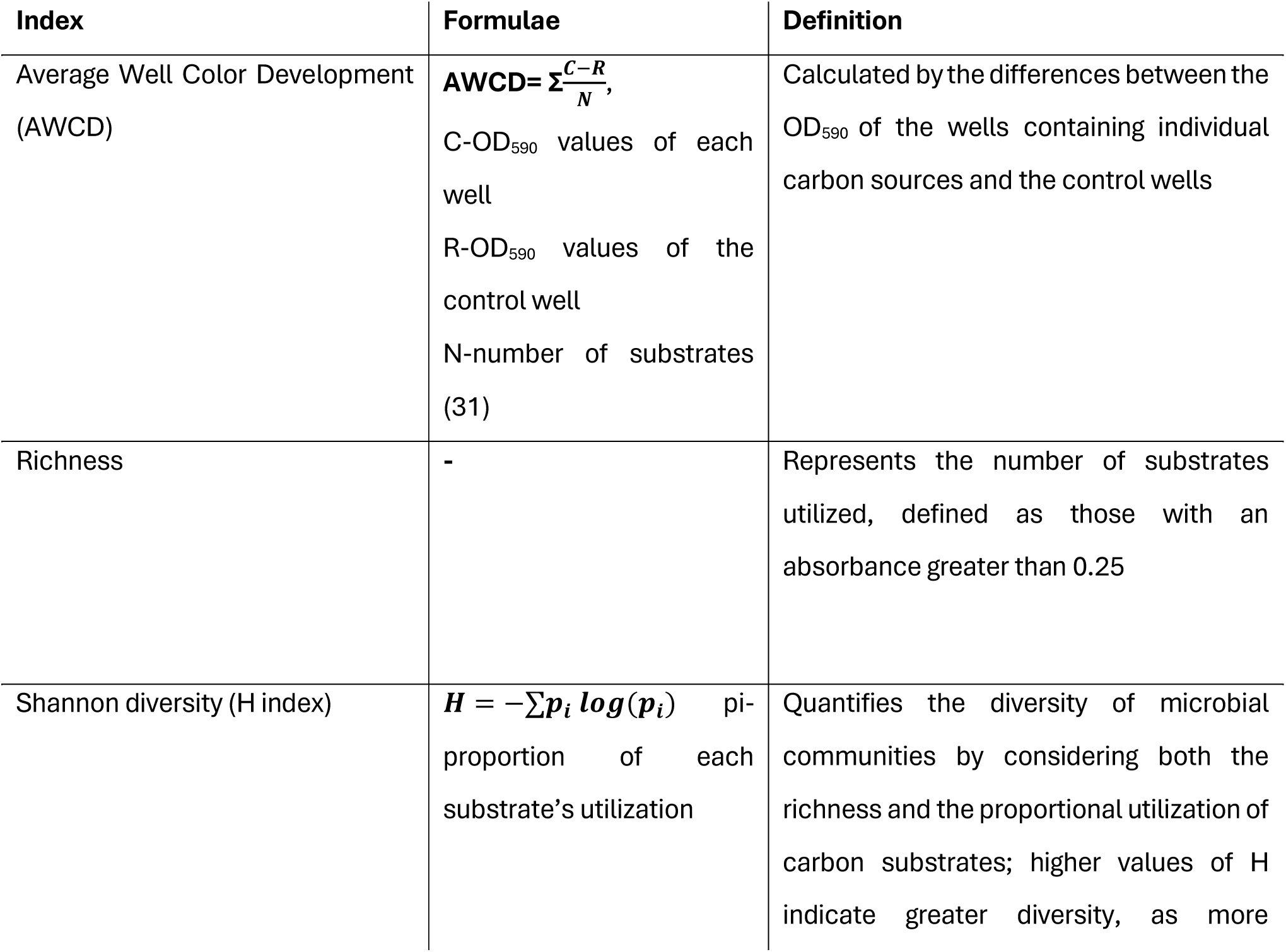
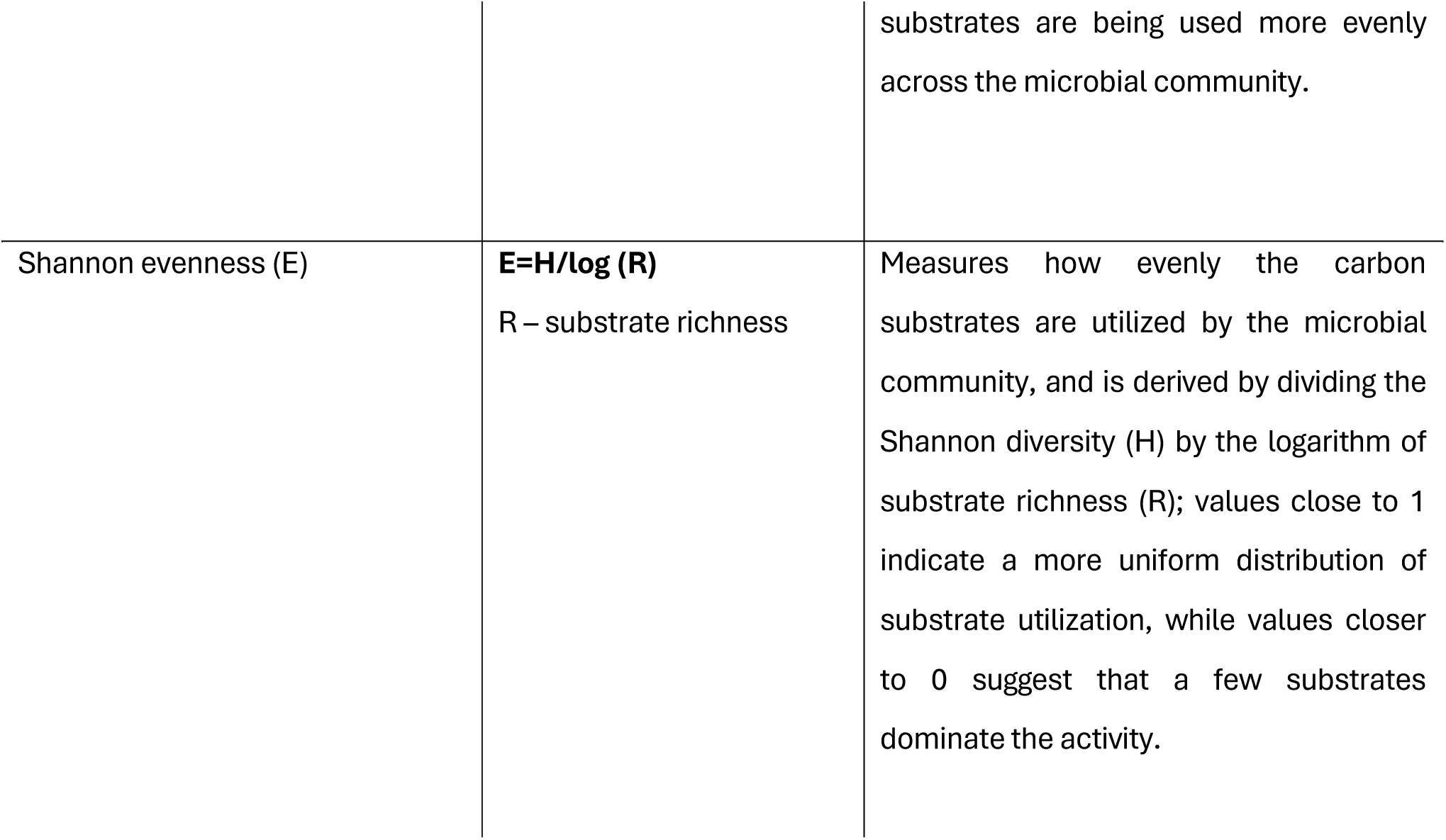
Microbial metabolic diversity indices and formulae used for calculations.

## Statistical analyses

### Generalized Additive Models (GAMs)

Generalized Additive Models (Simpson, 2024) were employed to explore the link between microbial diversity and geochemical factors. GAMs allow for non-linear relationships between predictor variables (e.g. values of geochemical or physico-chemical parameters) and response variables (e.g. microbial metabolic activity richness or diversity metrics). These models were fitted using the **mgcy** package, which applies smooth functions to each predictor. The microbial metrics, including richness, Shannon Diversity, and Shannon Evenness, were modeled as response variables with smooth terms for key geochemical predictors: pH, EC, and the concentrations of N, C, Na, Mg, K, Ca, P (Equation 1).

**Equation 1**. The formula for the GAMs which was applies smooth functions to each predictor.

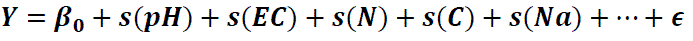

Where:

- *Y* is the microbial metric (Richness, Shannon Diversity, or Evenness),
- *s*(⋅) denotes a smooth function applied to each geochemical predictor.

These models allowed us to capture non-linear effects and interactions between environmental conditions and microbial diversity. For example, the smooth effect of pH might reveal an optimal pH range for microbial metabolic activity, while concentration of certain elements (e.g., Na and K) could influence specific functional groups in the microbial community. Moreover, the GAMs were visually inspected for their goodness of fit using residual diagnostics. Model checks and plots of the smooth functions for each predictor were generated to assess the relationship between diversity and geochemical factors.

### Principal Component Analysis (PCA)

The Principal Component Analysis (PCA) was used to investigate the joint variation in microbial community metabolic metrics and environmental geochemistry, by reducing the dimensionality of the dataset by identifying the principal components (PCs) that capture the majority of the variation in the datasets. Particularly, this is a powerful tool for identifying correlations between microbial metabolism (community-physiological profiles) and geochemical properties. In this case, PCA was applied to the combined dataset of microbial metrics such as AWCD, richness, Shannon Diversity, Shannon Evenness and physico-chemical variables (pH, EC, N, C, Na, Mg, K, Ca, P) using the **factoextra** package. These variables were scaled to ensure the comparability between measurements with different units for which the relationship between the principal components and the original standardized variables was established with the equation presented below (Equation 2), in which, Z represents the matrix of principal components, W is the weight matrix that defines the linear combination of the original variables, and X denotes the matrix of original standardized variables which includes microbial metrics and chemical parameters.

**Equation 2**. The formula used to reduce the dimensionality of the dataset while capturing the most significant variance, facilitating the interpretation of underlying patterns in microbial and chemical interactions.

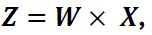

Biplots were created to visualize the relationship between microbial metrics and geochemical factors, displaying the sample clustering and contribution of variables.

### Data visualization

For the visualization of the results several graphs and plots were generated using **ggplot2** and **pheatmap** packages. Boxplots, line plots and scatterplots were used for the representation of microbial activity (AWCD and diversity metrics) across sample types and time points. Heatmaps coupled with hierarchical clustering were created to display metabolic activity patterns across carbon sources, samples and hours. The PCA biplot was used to explore the relationship between microbial metrics and geochemical factors. At the same time, the results of the GAMs depict the non-linear trend between chemistry and microbial diversity.

## Results

### Physico-chemical characteristics of cave water and sediment samples

Physico-chemical parameters, including pH, EC and the concentration of major elements (N, C, Na, Mg, K, Ca, P) were measured from sediment (limon and moonmilk included here) and water samples, with total nitrogen (TN) and total carbon (TC) considered for N, and C respectively (Table 3). Sediments show high calcium (Ca) concentrations, particularly in LAB3 (454,682 mg/kg) and LAB2 (226,206 mg/kg), reflecting the limestone-rich environment. The phosphorus (P) levels were highly variable, with some sediments, such as PMDF11 (41167 mg/kg) showing higher concentrations compared to PFDF2MM (104 mg/kg). On the other hand, in water samples, the electrical conductivity (EC) was notably higher, specifically PFDF4-APA showing the highest value (359 µS/cm). Overall, the pH of all the samples is alkaline with variations from 7.8 in sediment sample GHICA2 to 9.3 in LAB1 (sediment) and PFDFMMC (moonmilk). Carbon concentrations varied widely in sediments, ranging from < 100 mg/kg (e.g., PTDF1, LAB1) to 98,400 mg/kg (PMDF2), highlighting localized carbon hotspots. In water, concentrations were more consistent, ranging from 17.3 mg/L to 50.0 mg/L, reflecting its role in transporting dissolved carbon species. Sediments consistently exhibited higher concentrations of elements compared to water, indicating their function as nutrient and mineral reservoirs. The geochemical data were merged with the microbial metabolic diversity metrics by sample ID, allowing for a combined analysis of microbial community structure and environmental chemistry. However, certain elements (Al, Fe, S, Mn, As, Cr, Co, Ni, Cu, Zn) and water-specific parameters (TN, DC, DIC, DOC, SO₄^2-^, NO₃^−^, Cl^−^, PO₄^3-^) were excluded from the PCA due to either their low concentrations or the inability to integrate both sediment and water sample data in the same PCA. Nevertheless, aluminum (Al) and iron (Fe) dominated the sediment samples, with maximum concentrations of 63,089 mg/kg and 55,572 mg/kg, respectively. At the same time, sulfur (S) and trace metals like copper (Cu) and zinc (Zn) showed localized enrichment. In contrast, water samples exhibited significantly lower and more uniform concentrations, with aluminum peaking at 7.84 µg/L and sulfate reaching 48.6 mg/L, reflecting dilution effects and contrasting geochemical behaviors between sediments and water (supplementary table Table S2).

**Table 3.**
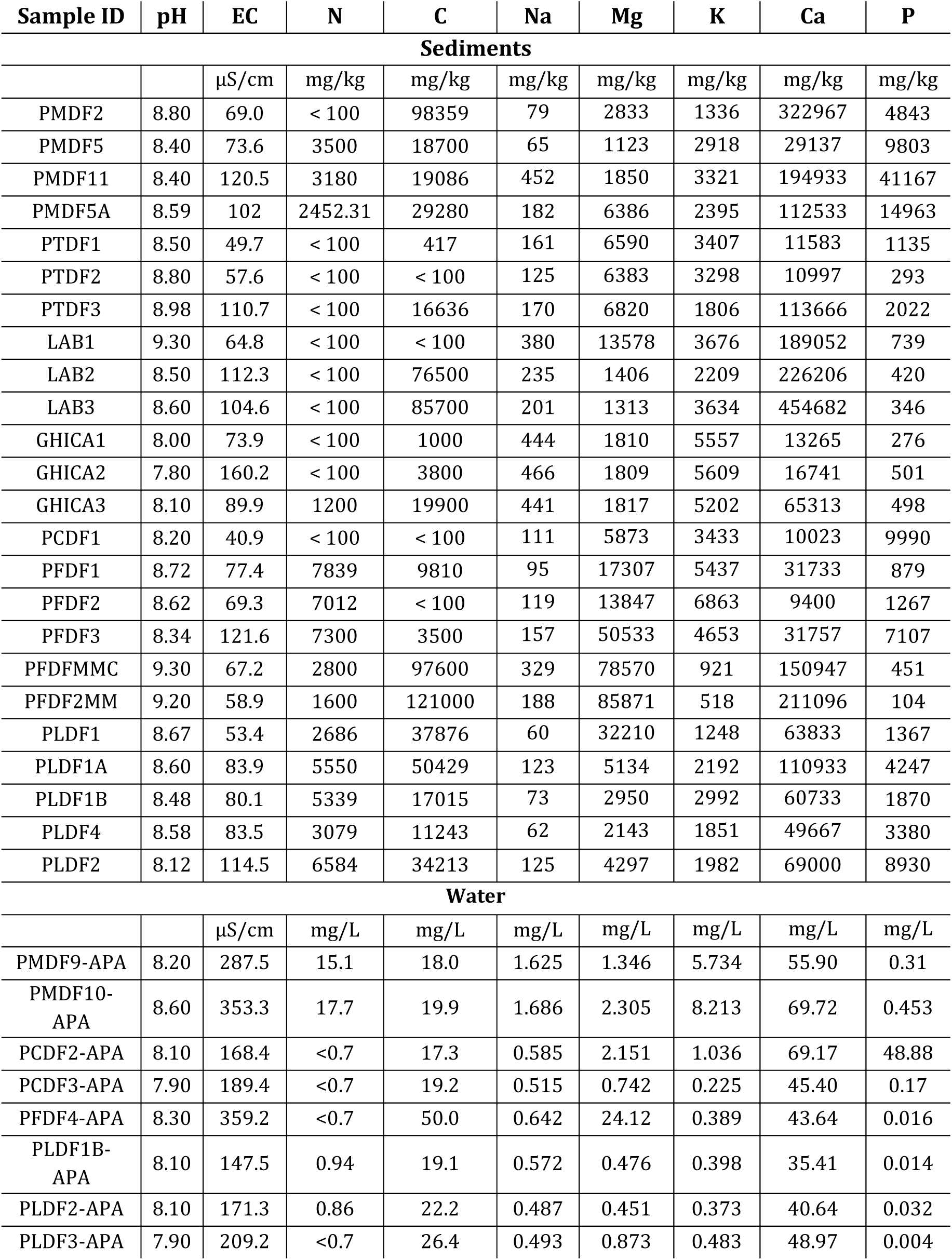
Physico-chemical parameters measured from sediment (including limon and moonmilk) and water samples, including pH, electrical conductivity (EC), and the concentrations of major elements (N, C, Na, Mg, K, Ca, P). Total nitrogen (TN) was considered as N, and total carbon (TC) as C, for water samples.

| Sample ID | pH | EC | N | C | Na | Mg | K | Ca | P |
| --- | --- | --- | --- | --- | --- | --- | --- | --- | --- |
| **Sediments** |  |  |  |  |  |  |  |  |  |
|  |  | μS/cm | mg/kg | mg/kg | mg/kg | mg/kg | mg/kg | mg/kg | mg/kg |
| PMDF2 | 8.80 | 69.0 | < 100 | 98359 | 79 | 2833 | 1336 | 322967 | 4843 |
| PMDF5 | 8.40 | 73.6 | 3500 | 18700 | 65 | 1123 | 2918 | 29137 | 9803 |
| PMDF11 | 8.40 | 120.5 | 3180 | 19086 | 452 | 1850 | 3321 | 194933 | 41167 |
| PMDF5A | 8.59 | 102 | 2452.31 | 29280 | 182 | 6386 | 2395 | 112533 | 14963 |
| PTDF1 | 8.50 | 49.7 | < 100 | 417 | 161 | 6590 | 3407 | 11583 | 1135 |
| PTDF2 | 8.80 | 57.6 | < 100 | < 100 | 125 | 6383 | 3298 | 10997 | 293 |
| PTDF3 | 8.98 | 110.7 | < 100 | 16636 | 170 | 6820 | 1806 | 113666 | 2022 |
| LAB1 | 9.30 | 64.8 | < 100 | < 100 | 380 | 13578 | 3676 | 189052 | 739 |
| LAB2 | 8.50 | 112.3 | < 100 | 76500 | 235 | 1406 | 2209 | 226206 | 420 |
| LAB3 | 8.60 | 104.6 | < 100 | 85700 | 201 | 1313 | 3634 | 454682 | 346 |
| GHICA1 | 8.00 | 73.9 | < 100 | 1000 | 444 | 1810 | 5557 | 13265 | 276 |
| GHICA2 | 7.80 | 160.2 | < 100 | 3800 | 466 | 1809 | 5609 | 16741 | 501 |
| GHICA3 | 8.10 | 89.9 | 1200 | 19900 | 441 | 1817 | 5202 | 65313 | 498 |
| PCDF1 | 8.20 | 40.9 | < 100 | < 100 | 111 | 5873 | 3433 | 10023 | 9990 |
| PFDF1 | 8.72 | 77.4 | 7839 | 9810 | 95 | 17307 | 5437 | 31733 | 879 |
| PFDF2 | 8.62 | 69.3 | 7012 | < 100 | 119 | 13847 | 6863 | 9400 | 1267 |
| PFDF3 | 8.34 | 121.6 | 7300 | 3500 | 157 | 50533 | 4653 | 31757 | 7107 |
| PFDFMMC | 9.30 | 67.2 | 2800 | 97600 | 329 | 78570 | 921 | 150947 | 451 |
| PFDF2MM | 9.20 | 58.9 | 1600 | 121000 | 188 | 85871 | 518 | 211096 | 104 |
| PLDF1 | 8.67 | 53.4 | 2686 | 37876 | 60 | 32210 | 1248 | 63833 | 1367 |
| PLDF1A | 8.60 | 83.9 | 5550 | 50429 | 123 | 5134 | 2192 | 110933 | 4247 |
| PLDF1B | 8.48 | 80.1 | 5339 | 17015 | 73 | 2950 | 2992 | 60733 | 1870 |
| PLDF4 | 8.58 | 83.5 | 3079 | 11243 | 62 | 2143 | 1851 | 49667 | 3380 |
| PLDF2 | 8.12 | 114.5 | 6584 | 34213 | 125 | 4297 | 1982 | 69000 | 8930 |
| **Water** |  |  |  |  |  |  |  |  |  |
|  |  | μS/cm | mg/L | mg/L | mg/L | mg/L | mg/L | mg/L | mg/L |
| PMDF9-APA | 8.20 | 287.5 | 15.1 | 18.0 | 1.625 | 1.346 | 5.734 | 55.90 | 0.31 |
| PMDF10-APA | 8.60 | 353.3 | 17.7 | 19.9 | 1.686 | 2.305 | 8.213 | 69.72 | 0.453 |
| PCDF2-APA | 8.10 | 168.4 | <0.7 | 17.3 | 0.585 | 2.151 | 1.036 | 69.17 | 48.88 |
| PCDF3-APA | 7.90 | 189.4 | <0.7 | 19.2 | 0.515 | 0.742 | 0.225 | 45.40 | 0.17 |
| PFDF4-APA | 8.30 | 359.2 | <0.7 | 50.0 | 0.642 | 24.12 | 0.389 | 43.64 | 0.016 |
| PLDF1B-APA | 8.10 | 147.5 | 0.94 | 19.1 | 0.572 | 0.476 | 0.398 | 35.41 | 0.014 |
| PLDF2-APA | 8.10 | 171.3 | 0.86 | 22.2 | 0.487 | 0.451 | 0.373 | 40.64 | 0.032 |
| PLDF3-APA | 7.90 | 209.2 | <0.7 | 26.4 | 0.493 | 0.873 | 0.483 | 48.97 | 0.004 |

### Patterns of carbon source utilization in cave samples

The carbon substrate metabolization patterns significantly varied across all assayed cave samples (Figure 2). A higher OD_590_ was recorded for the samples inoculated in the wells containing Tween 40, putrescine and D-xylose, indicating a higher rate of microbial activity toward degradation of these compounds. Conversely, lower OD_590_ values after substrate consumption were exhibited in the wells containing 2-hydroxy benzoic acid and L-threonine suggesting limited microbial capacity to metabolize these compounds. The more easily accessible compounds, such as pyruvic acid methyl ester and D-galacturonic acid showed consistently higher activity, highlighting the presence of generalist metabolic traits.

**Figure 2.**
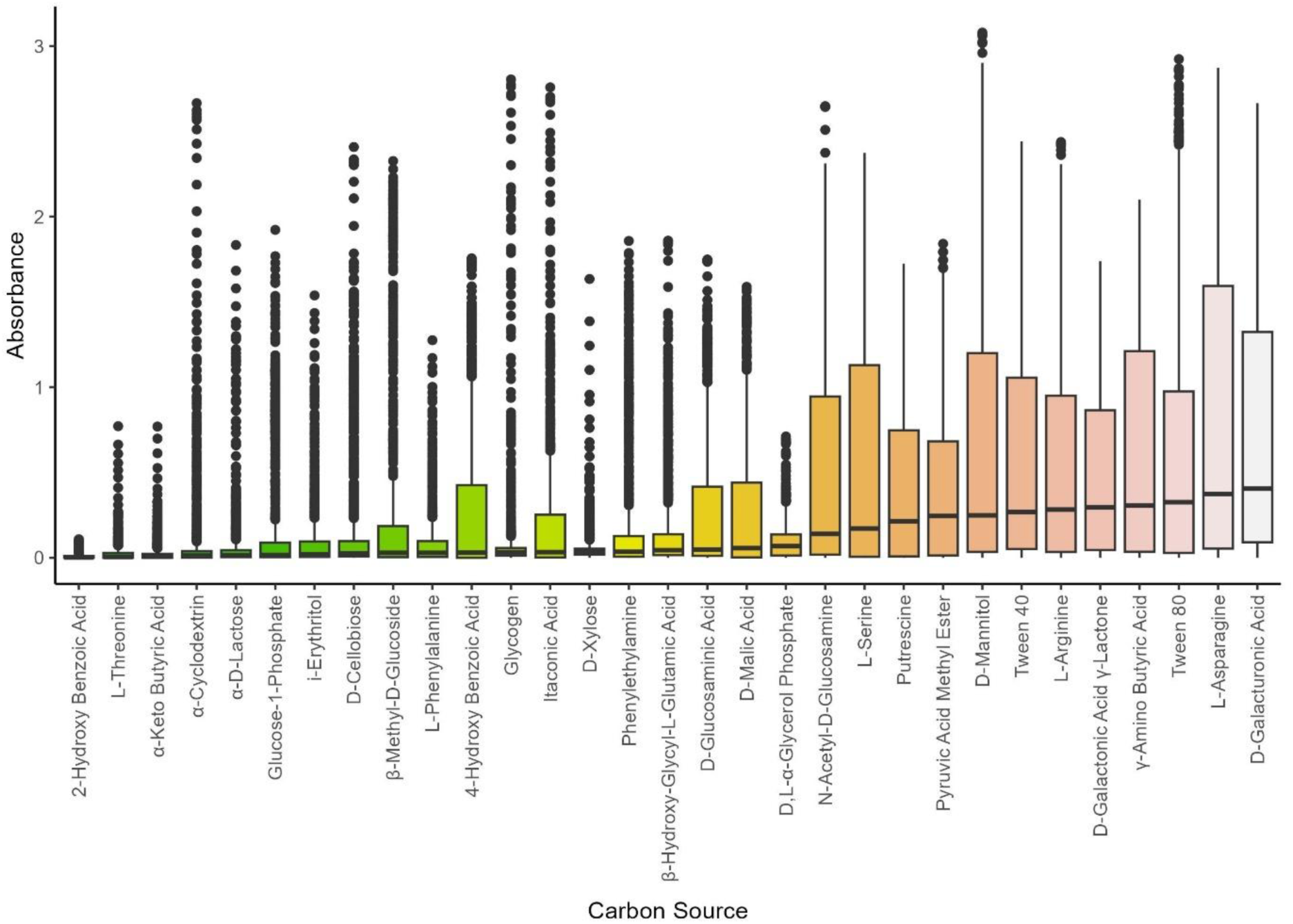
Box plot showing the median absorbance for the metabolization of 31 carbon substrates by microbial communities, highlighting differences in substrate utilization and associated metabolic activity. Each box represents the interquartile range (IQR), with whiskers extending to 1.5 times the IQR, and individual dots showing outlier data points. The color coding differentiates the substrates but does not carry a specific meaning. Excepting that, in Table 4 are summarized the average absorbances for selected carbon substrates for which the microbial communities from water samples exhibited the highest metabolic activity across several carbon sources.

**Figure 3.**
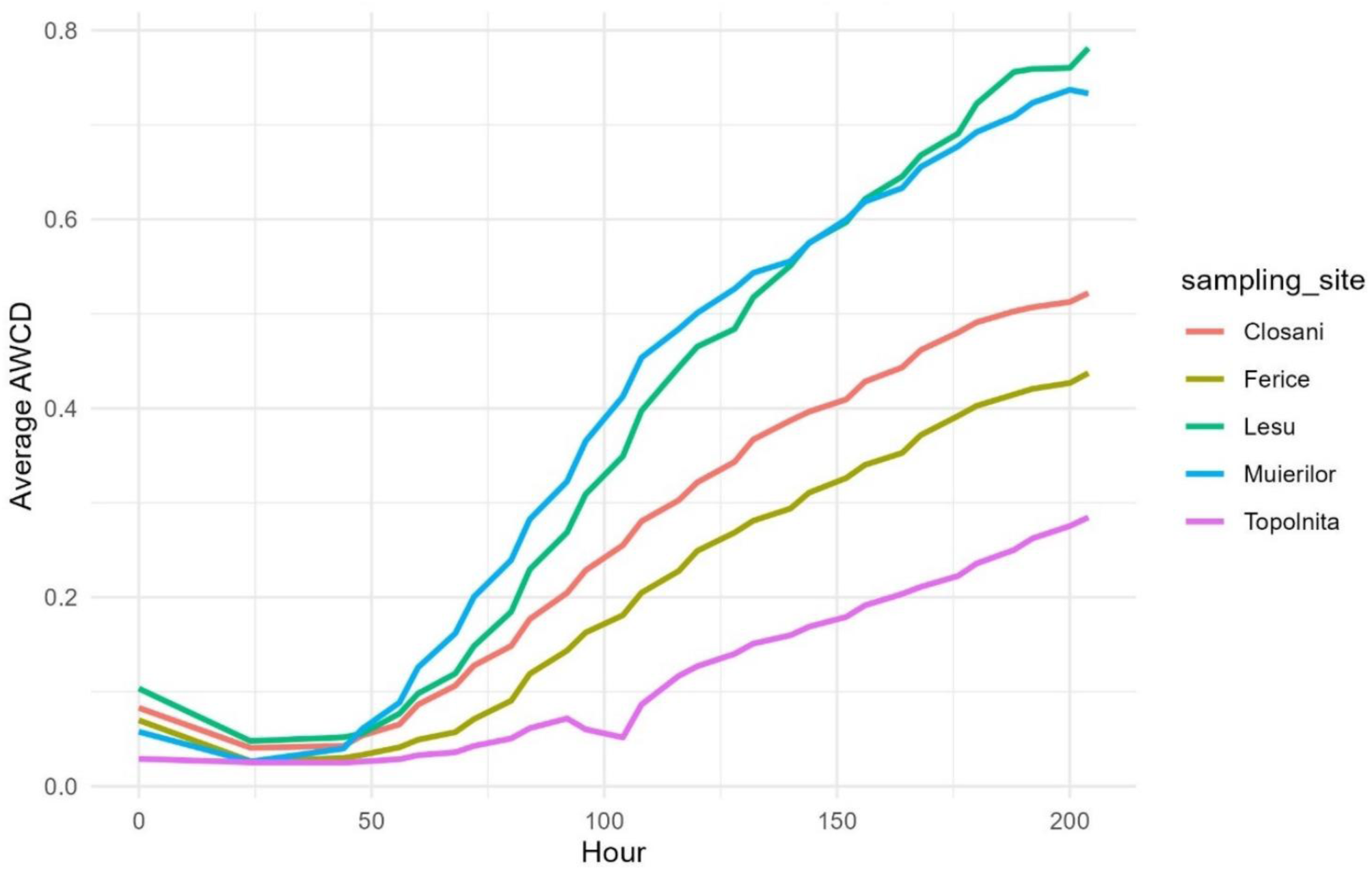
Average-well Color Development (AWCD) over time for all samples across five sampling sites: Closani, Ferice, Lesu, Muierilor, and Topolnița. The trends highlight site-specific differences in substrate utilization rates, with samples from Muierilor showing the highest activity, followed by Lesu, Closani, Ferice, and Topolnița. Figure 3 shows the AWCD values over time for all samples across the sampling sites, whereas Figure 4 and Table S1 shows the AWCD values over time by sample type reflecting the overall metabolic activity from each environment. By far, the water samples have an average-well color development value rising sharply after 50 hours of incubation and a continuous rapid increase, reaching a final value of 0.8, thus highlighting the highest overall metabolic activity among sample types. Sediment samples showed a moderate AWCD which increased steadily over time with approximately 0.5 at the end of the incubation period. This trend suggested an active heterotrophic microbial community capable of utilizing various carbon substrates, yet at a reduced rate, potentially affected by sediment type, moisture content, or microbial composition at the sampling locations. Samples from moonmilk deposits demonstrated the lowest AWCD, showing a gradual and minimal increase during the incubation period, reflecting a less metabolically diverse microbial community. The overall AWCD patterns across the sample types reflect significant differences in microbial diversity and functional capabilities across these distinct cave environments.

**Table 4.** Average absorbance (OD_590_) for selected carbon sources (± Standard Deviation)

| Carbon Source | Water<br>(Mean ± SD) | Sediment<br>(Mean ± SD) | Moonmilk<br>(Mean ± SD) |
| --- | --- | --- | --- |
| D-mannitol | 1.05 ± 0.85 | 0.63 ± 0.82 | 0.05 ± 0.03 |
| N-acetyl-D-glucosamine | 1.03 ± 0.72 | 0.42 ± 0.60 | 0.02 ± 0.01 |
| L-serine | 0.80 ± 0.68 | 0.55 ± 0.68 | 0.00 ± 0.00 |
| Tween 80 | 1.03 ± 0.79 | 0.49 ± 0.60 | 0.20 ± 0.24 |
| D-galacturonic acid | 1.22 ± 0.75 | 0.60 ± 0.63 | 0.13 ± 0.13 |

The average absorbances for selected carbon substrates indicated that the microbial community from the water samples exhibited the highest metabolic activity for a wide range of carbon sources (Table 4). D-galacturonic acid (mean absorbance = 1.22 ± 0.75), D-mannitol (1.05 ± 0.85), N-acetyl-D-glucosamine (1.03 ± 0.72) and Tween 80 (1.03 ± 0.79). In contrast, the microbial communities from the moonmilk samples showed the lowest metabolic response, with absorbance values remaining below 0.5 for most carbon sources.

We evaluated the temporal dynamics based on median absorbance values for the carbon source utilization across the sample types (Figure 2). In water samples an overall higher C substrate utilization compared to the other samples was noted, specifically for substrates such as D-galacturonic acid, pyruvic acid methyl ester, Tween 40 and N-acetyl-D-glucosamine (Table 4). Certain C-sources, like putrescine and L-arginine, represent substrates for bacterial enzymatic activity, indicating similar patterns in sediment and water, but with generally higher utilization by water-associated communities. The moonmilk microbial communities overall show the lowest C-substrate utilization with many substrates exhibiting little to no increase in absorbance over time (e.g., γ-amino butyric acid, β-methyl-D-glucoside, D-cellobiose), except for D-xylose and 2-hydroxy benzoic acid which depict a higher median absorbance compared to sediment and water samples.

Similarly, the temporal dynamics of median absorbances across the carbon substrates were assessed for sampling sites. Muierilor Cave is the site which consistently shows the highest microbial-driven metabolic activity across the majority of substrates with elevated absorbance values for D-galacturonic acid, L-serine, D-malic acid, L-asparagine, and D-mannitol, suggesting that the communities from this sampling site are capable of utilizing a wide variety of substrates. On the other hand, the microbial communities associated to samples from Topolnița Cave generally show the lowest activity, except for 2-hydroxy benzoic acid. Also, separate graphs were created for each sample type (Supplementary Figure S2 – S4) which helps to have an overview of differences between samples and the capabilities of the communities harboring those sampling points. For example, the substrate metabolization by the microbial communities of the water samples collected from Cloșani exhibits a higher OD value (Supplementary Figure S2) for 19 carbon sources in water making it one of the most active sites for this sample type. In contrast, the utilization of C-sources by the microbial communities among water samples from Muierilor shows a lower activity with a few exceptions for substrates like 2-hydroxy benzoic acid, D-malic acid, D-xylose, L-arginine, L-asparagine, Tween 40 and Tween 80. The sediment samples (Supplementary Figure S3) show almost similar activity in samples from Leșu and Muierilor, yet higher in Leșu, whereas Cloșani, Ferice and Topolnița depict approximately similar lower activity except for 2-hydroxy benzoic acid (Topolnița), α-Keto-Butyric Acid (Cloșani) and L-serine (Ferice). In the moonmilk samples from a single sampling point (Ferice; Supplementary Figure S4), the median absorbance revealed that Tween 80 was the fastest consumed substrate followed by Tween 40, and L-phenylalanine.

### Estimation of overall carbon metabolization by Average-Well Color Development (AWCD)

The average-well color development (AWCD) at each sampling point during 200 hours shows the carbon substrates being utilized and their rate of consumption, providing clues on overall metabolic activity. The overall trend for all sampling locations indicated that substrate utilization typically increases after 50 hours of plate incubation. The samples from the show cave Muierilor showed the most significant increase between 50 and 120 hours, whereas samples from Leșu exhibited a comparable sharp rise ultimately achieving a final AWCD value of 0.8 at the end of the experiment. Cloșani samples exhibited moderate activity with a steadier utilization pattern over time, while samples from Ferice displayed gradual increase in activity but lower AWCD values, indicating a community that is either less metabolically diverse or less active. Samples from Topolnița Cave showed limited activity, as indicated by a gradual, flat curve in the early hours, possibly due to metabolic constraints or geochemical conditions.

### Microbial metabolic diversity metrics across cave samples

Temporal dynamics of metabolic richness and diversity

The Shannon diversity metrics shown in Figure 5 boxplot estimate the microbial diversity for each sampling location. These results are supplemented by the findings summarized in Table 5 which shows the maximum richness and H index by sample type, along with the supplementary Figure S7 which plots the Shannon diversity by sample type. Water samples showed the highest median value, close to 3.0, indicating that the communities present here are the most metabolically diverse and evenly distributed substrate consumption. In contrast, sediment samples exhibited moderate substrate consumption diversity, while the moonmilk samples have the lowest median substrate metabolization suggesting the presence of a less diverse community with limited substrate preferences (Supplementary Figure S7). These findings are also reflected in the high median diversity values for sites like Cloșani and Ferice (Figure 4), which might indicate that both water and sediment samples from Cloșani support the most metabolically diverse microbial communities. Despite the higher values of these indices registered for Ferice Cave, the moonmilk samples collected from this site depict the lowest richness and diversity (Table S1 and Supplementary Figure S7), possibly due to a more specialized or less diverse community. The sampling locations from Leșu and Muierilor show moderate diversity with a median value below 2.7 (Figure 4) with a higher C-substrate metabolization diversity in sediments compared to water (Table S1). The least and lowest Shannon diversity values (around 2.5) is found in samples from Topolnița. Comparing the both graphs it is clearer that sampling location and sample type are the key factors influencing microbial metabolic diversity, with water samples showing the highest complexity.

**Figure 4.**
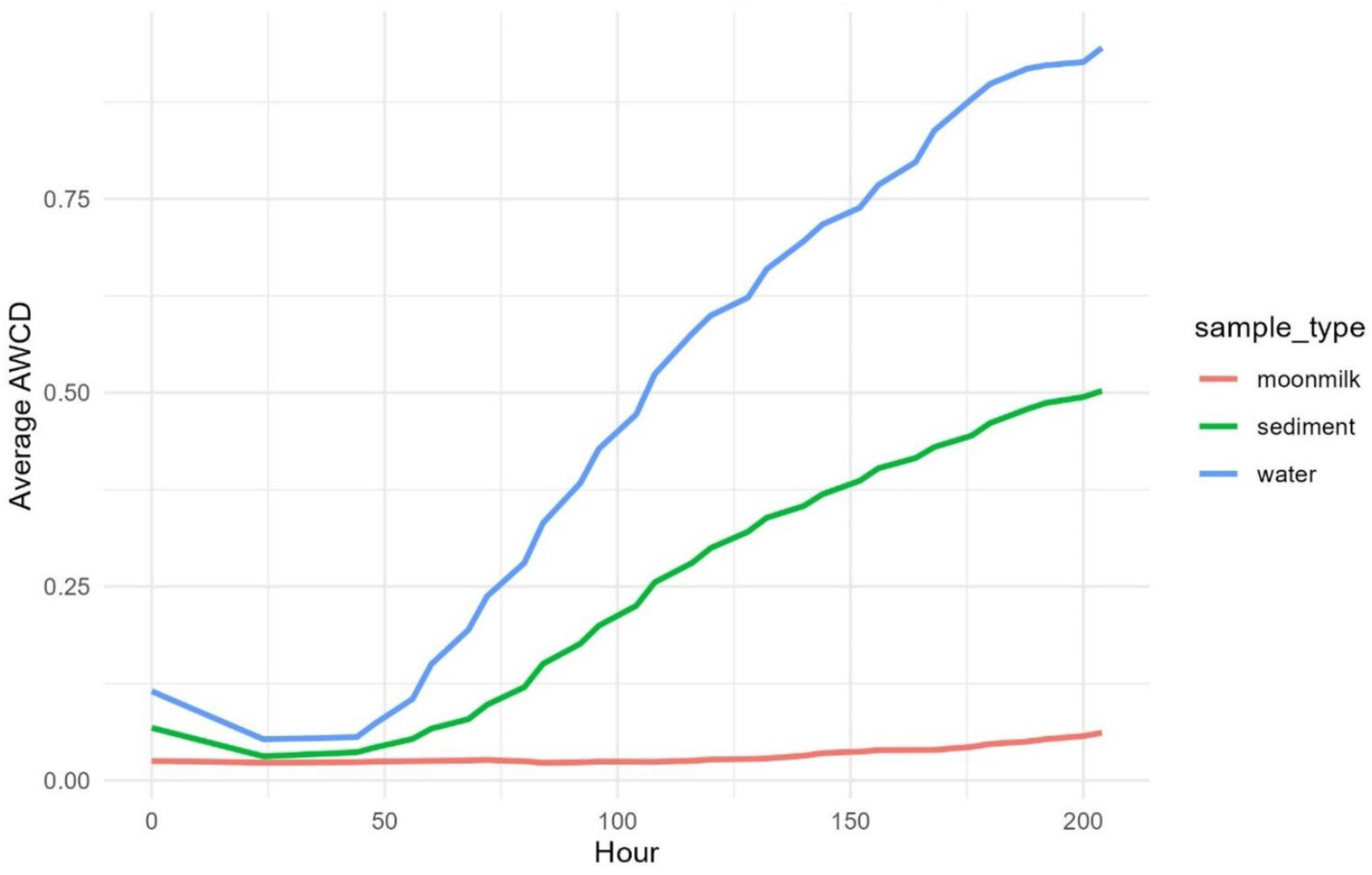
Average-well Color Development (AWCD) over time for microbial communities from three sample types: moonmilk, sediment, and water.

**Figure 5.**
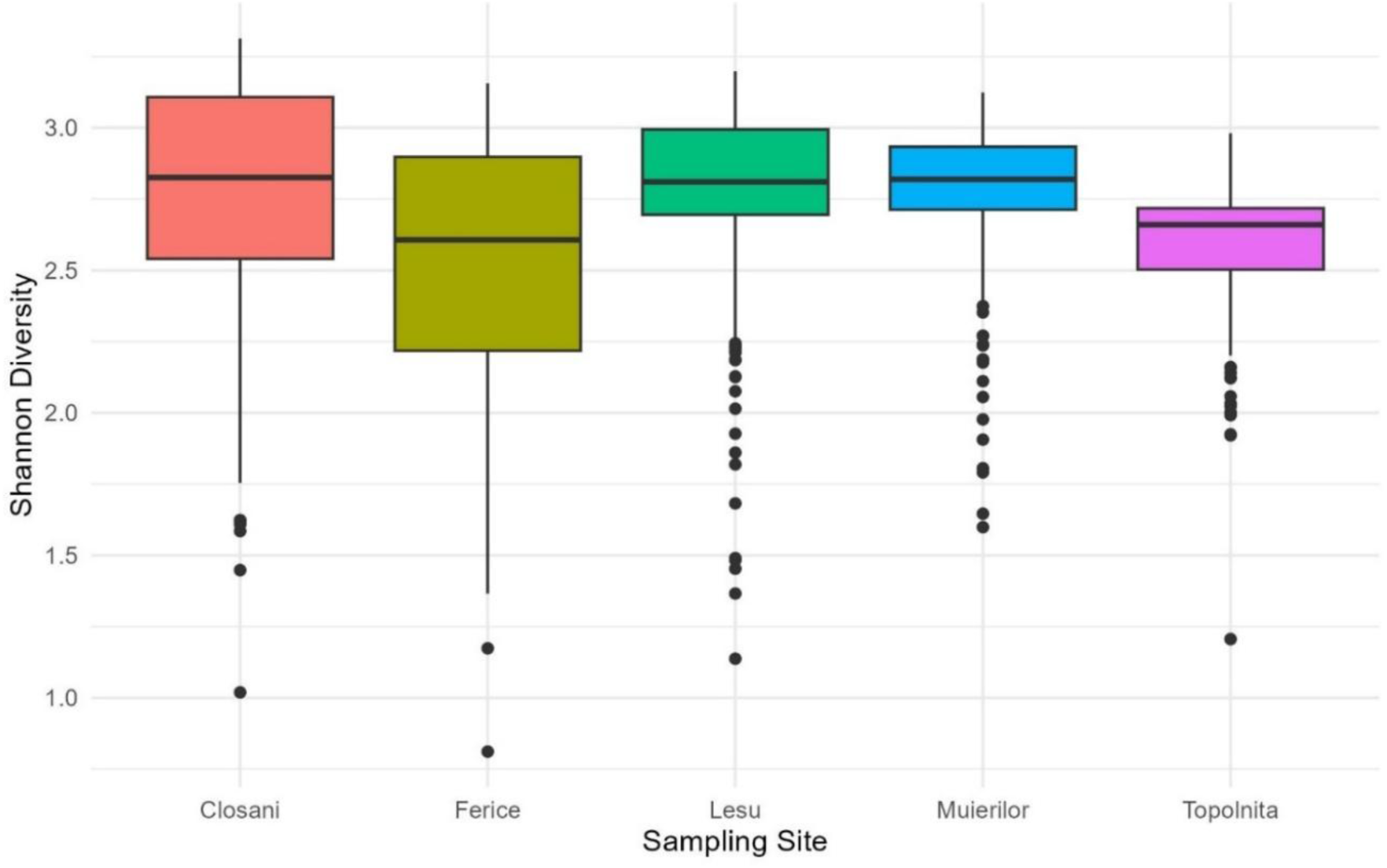
Box plot of Shannon diversity indices based on substrate utilization patterns for microbial communities across sampling sites (Closani, Ferice, Lesu, Muierilor, and Topolnița). The box colors correspond to different sampling sites, while the black dots represent outliers in Shannon diversity values.

**Table 5.** Maximum richness and Shannon diversity index by sample type across sampling sites. Richness represents the total number of substrates metabolized, while the Shannon diversity index reflects both the richness and evenness of substrate utilization.

| Sample Type | Max Richness (Water) | Max Richness (Sediment) | Max Richness (Moonmilk) | Shannon Diversity (Water) | Shannon Diversity (Sediment) | Shannon Diversity (Moonmilk) |
| --- | --- | --- | --- | --- | --- | --- |
| Closani | 54 | 76 | - | 6.45 | 26.98 | - |
| Ferice | 24 | 78 | 4 | 3.16 | 14.28 | 6.13 |
| Lesu | 56 | 101 | - | 8.64 | 14.94 | - |
| Muierilor | 49 | 62 | - | 6.19 | 11.84 | - |
| Topolnita | - | 29 | - | - | 8.62 | - |

### Correlation of the AWCD and richness values

A positive correlation was observed between AWCD (metabolic activity) and richness (substrate utilization diversity) across all sample types. Despite some site-specific variations, evidence suggests that as AWCD increases, richness also tends to increase.

The graphical representation for each sampling site or sample category (Figure 5 and Supplementary Figure S8) presents the time series of AWCD and richness values associated with each sample type. All measurements at these time points have been included in the final scatter plots, illustrating the temporal variability in microbial metabolic activity and substrate utilization diversity.

The results (Figure 5) indicated a heterogeneity among sampling sites, suggesting that Cloșani may support communities with greater metabolic versatility and diversity of consumed substrates. An overlap in richness values across different sites when AWCD values are low (0.0 – 0.5) might indicate that multiple sampling sites have similar species richness despite having different metabolic activities. For AWCD values between 0.5 – 1.0, richness increased more consistently among sites (e.g., Leșu, Muierilor, Ferice) and a broader spread can be seen. Interestingly, Cloșani had a wider spread in terms of both AWCD and richness with a positive trend, but mostly extended to the highest AWCD values (1.0 – 1.5) and richness (close to 30). Also, a few samples from Ferice also showed a positive trend, but this site actually depicted an intermediate metabolic activity and species richness. Supplementary Figure S8 shows the widest range of richness and AWCD values for sediment samples, followed by water samples which encounter the highest richness values. With few data points and low values moonmilk samples remain with the lowest richness (< 10) and metabolic activity (< 0.1).

### Correlations between microbial metrics and chemistry

Correlation analysis (Figure S9 and Figure S10) revealed several key relationships between microbial activity metrics and environmental chemistry variables. AWCD was significantly positively correlated with sodium (Na) (r = 0.56, p < 0.01) and phosphorus (P) (r = 0.45, p < 0.05). Shannon diversity was strongly correlated with electrical conductivity (EC) (r = 0.49, p < 0.05) and magnesium (Mg) (r = 0.55, p < 0.05). Significant correlations indicate that these environmental parameters likely play an important role in shaping microbial capability to degrade different C-substrates and function in cave environments (Table 6).

**Table 6.** Pearson correlation coefficients for microbial activity metrics and environmental chemistry parameters. A single asterisk (*) denotes significance at the p < 0.05 level, and a double asterisk (**) denotes significance at the p < 0.01 level.

| Metric | Na | P | EC | Mg | K |
| --- | --- | --- | --- | --- | --- |
| AWCD | 0.56** | 0.45* | 0.39 | 0.55* | 0.37 |
| Richness | 0.44* | 0.32 | 0.48* | 0.49* | 0.30 |
| Shannon Diversity | 0.50* | 0.40 | 0.49* | 0.55* | 0.41* |

### Analysis of microbial activity and carbon utilization in cave sediments

The heatmaps complemented by hierarchical clustering dendrograms have been generated to evaluate the diversity of microbial metabolic activities across cave samples. The dendrograms facilitate understanding the biological links between sediment, water and moonmilk sample types. Hierarchical clustering dendrograms show differences in microbial communities’ metabolic profiles and composition across sample types.

The hierarchical clustering of AWCD by sample and time highlights four distinct clusters of microbial activity across different cave environments (Figure S11). Sediment samples from Cloșani and Muierilor show lower AWCD values across time, while water samples such as PCDF2-APA (Cloșani) and PLDF1B-APA (Leșu), form separate clusters with much higher AWCD values, suggesting more active microbial communities. Moonmilk samples (e.g., PFDF2MM and PFDFMMC) cluster along with most of the sediment samples, exhibiting lower levels of AWCD, which may indicate similar metabolic limitations or substrate availability.

Clustering of substrate richness by sample and time (Figure S11) offers valuable insights into the C-substrate utilization diversity of microbial communities, for which the highest substrate richness is observed in samples such as PCDF2-APA, PMDF9-APA and PMDF10-APA particularly at later time points (116-204), indicated by the color scale. In contrast, samples like PFDF1 to PFDFMMC have consistently low richness over time as shown by the dark purple shading. Temporal clustering indicates that samples from similar time points (e.g., 72–108 hours and 144–168 hours) often group together, highlighting periods of peak microbial activity and substrate diversity.

The Shannon diversity heatmap accounts for both substrate richness and evenness, shows clear separation of the sample types in the dendrogram (Figure S12). A higher Shannon diversity values is consistently observed in samples such as GHICA2, PCDF2-APA, PMDF9-APA and PMDF10-APA, particularly at time points from 100-204 hours. Samples such as PFDF2, GHICA1 and PLDF3-APA depict a reduced metabolic diversity for almost all time points, indicating low microbial evenness.

Moreover, a hierarchical clustering analysis was conducted to explore the grouping patterns of microbial activity based on the utilization of carbon sources over time (Figure S13). Additionally, the results highlight the clustering of individual samples (Figure S14) and the clustering of sample types (Figure S15) in relation to specific carbon source utilization patterns based on the median absorbance values. The key findings revealed that L-asparagine and D-galacturonic acid are associated highest metabolic activity at later points (over 120 hours), as indicated in the Figure S13. Tween 40 and Tween 80 exhibit increasing absorbances through mid-late time intervals (80-204 hours), indicating that these carbon sources are metabolized following a lag phase, possibly due to specific microbial community shifts (Figure S13).

For several carbon sources the metabolic activity peaks between 100-204 hours suggesting a period of intense microbial substrate utilization (e.g., D-galacturonic acid γ-lactone, L-arginine, L-serine).

Other carbon sources, including L-threonine, α-cyclodextrin, and α-keto butyric acid, exhibit minimal metabolic activity throughout the entire time period, as shown by the dark purple shading. L-asparagine, Tween 80, and D-galacturonic acid (Supplementary Figure S14) show high absorbance across multiple samples, indicating these carbon sources are widely utilized by diverse microbial communities. On the other hand, certain carbon sources such as L-threonine and α-keto butyric acid, demonstrate minimal utilization in the majority of samples, as evidenced by consistent dark purple shading, indicating a restricted microbial ability to metabolize these substrates. This hierarchical clustering (Supplementary Figure S14) reveals distinct groups of samples with similar metabolic activity patterns, indicating potential community structuring based on carbon source availability. In the Supplementary Figure S15 water samples exhibit the highest metabolic activity, particularly for carbon sources like L-asparagine and D-galacturonic acid as indicated by the yellow coloration. Samples of sediment and moonmilk exhibit diminished overall metabolic activity, with the majority of carbon sources presenting low absorbance values (blue/purple), indicating a more restricted range of substrate use relative to water.

### Environmental drivers of carbon substrate utilization

PCA was employed to examine the relationships between microbial metrics, including richness, Shannon diversity, and evenness along with several chemical variables such as pH, EC, N, C, Na, Mg, K, Ca, and P. The analysis promotes dimensionality reduction while preserving the most significant patterns within the dataset. Figure 8, illustrates the microbial metrics and chemical variables within a reduced component space. Moreover, the scatterplot matrix (Supplementary Figures S9 and S10) highlight the links between cave physico-chemical variables and microbial metrics.

**Figure 6.**
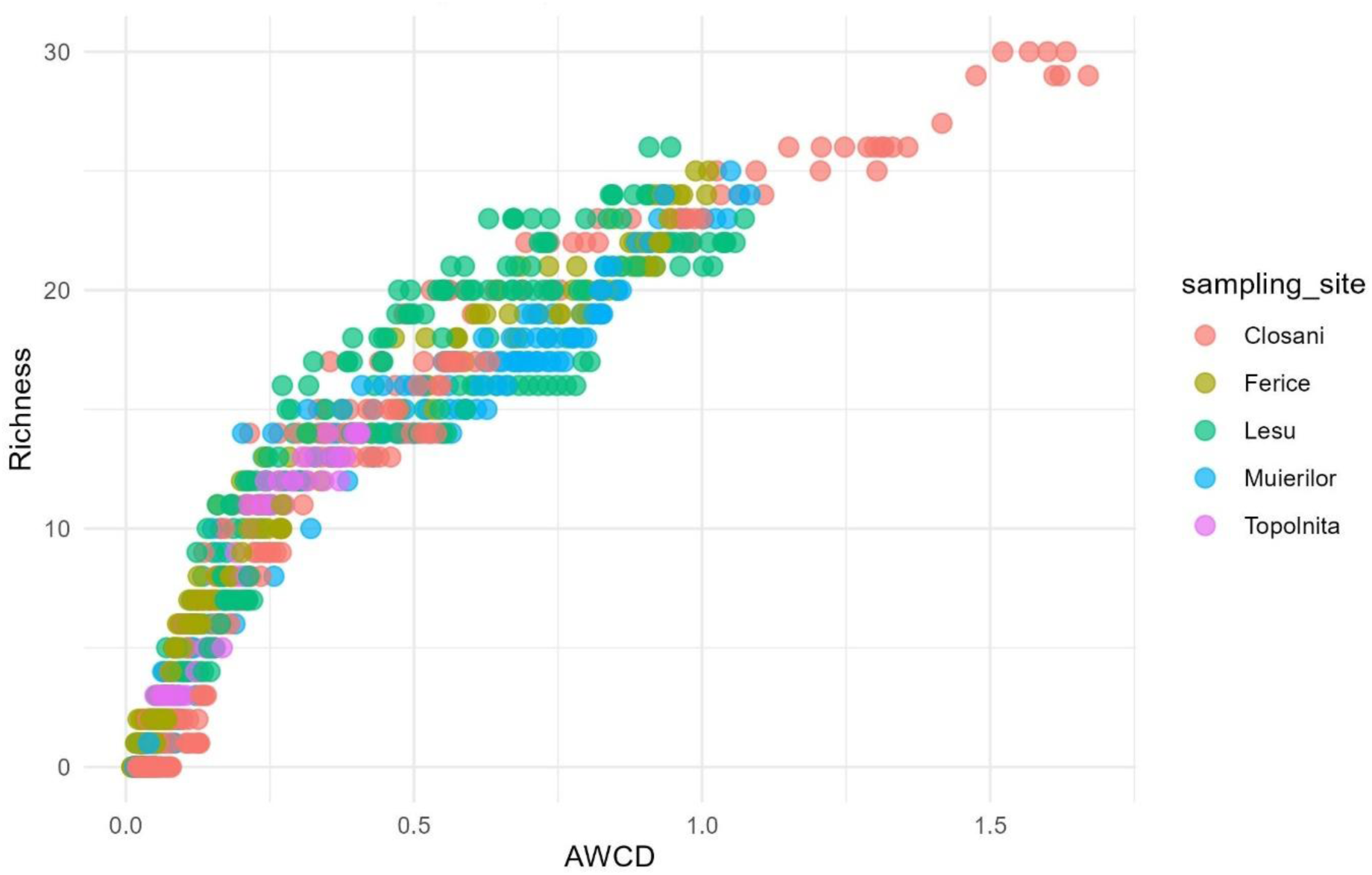
The scatter plot of AWCD and richness values measured at different time points over a total duration of 0-204 hours for each sample from each sampling site. All the measurements were included in the scatter plot to capture the temporal dynamics.

**Figure 7.**
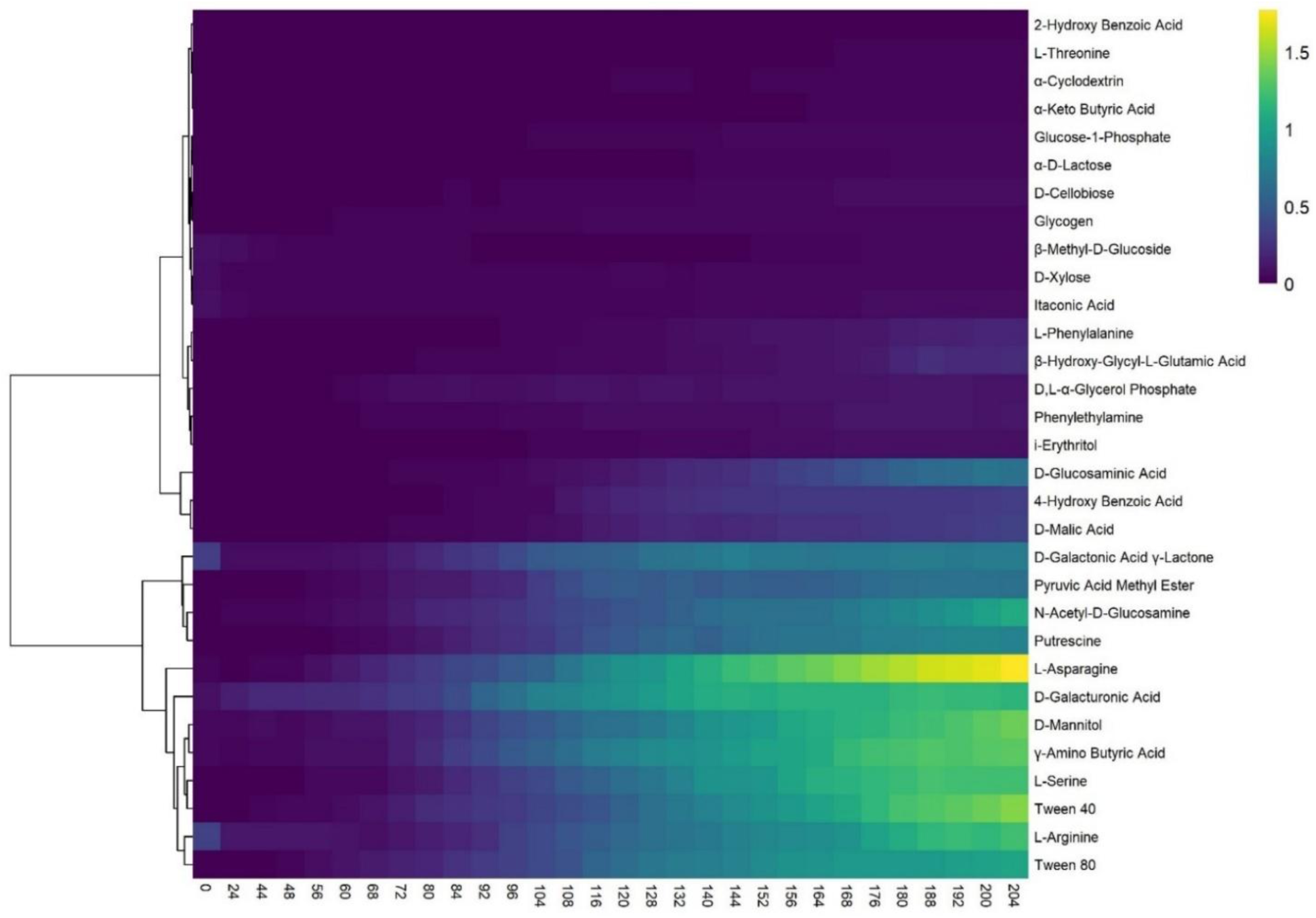
Heatmap plot of the median absorbance of microbial metabolic diversity across the carbon source types over time. The color scale represents absorbance values, with yellow indicating higher metabolic activity and purple indicating lower activity. Hierarchical clustering was applied to both carbon sources (rows) and time points (columns), revealing patterns of substrate utilization by the microbial communities over time.

**Figure 8.**
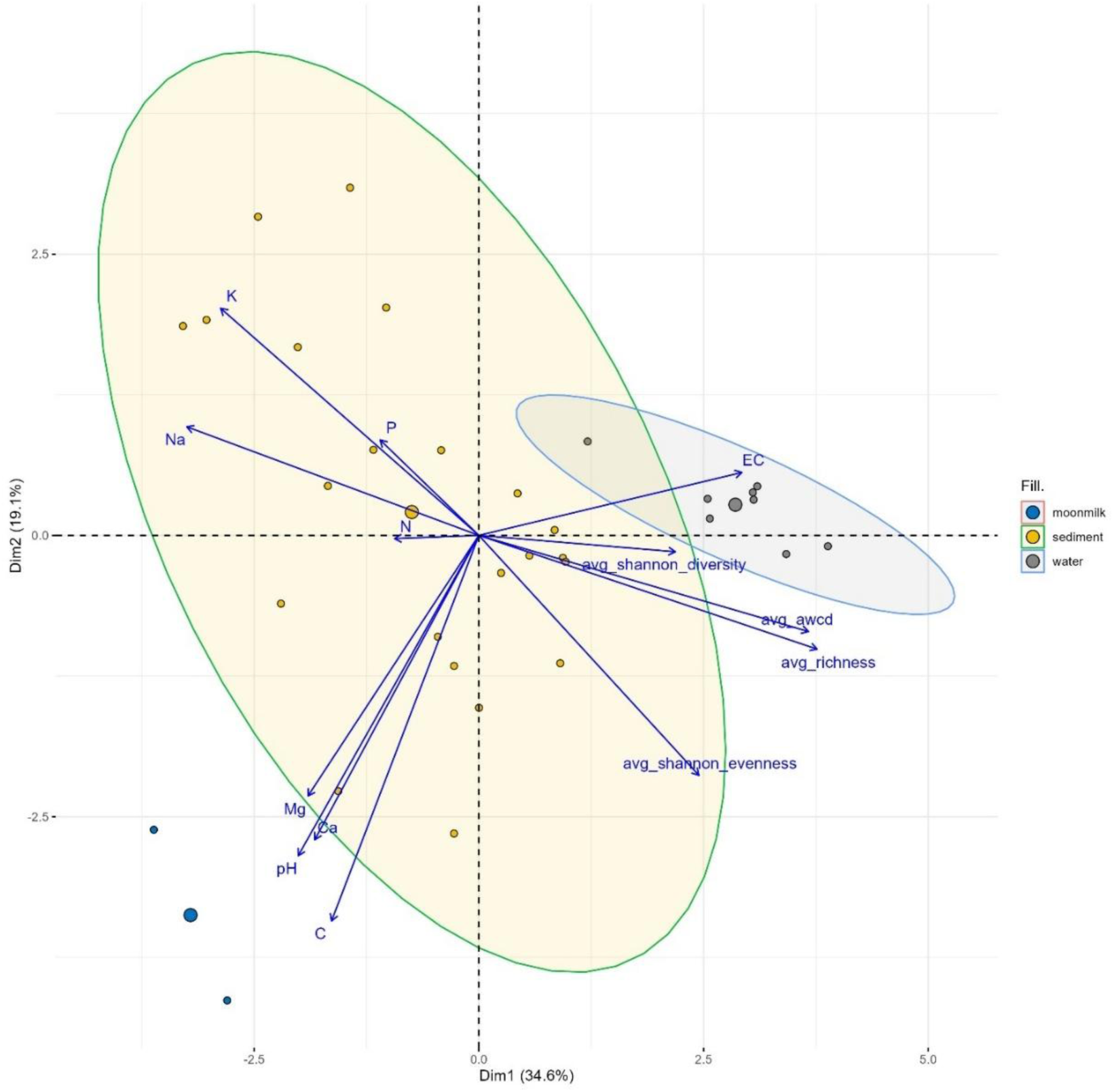
Principal Component Analysis (PCA) biplot showing the relationships between microbial metrics and cave chemistry variables across sample types. The first two principal components explain 34.6% and 19.1% of the variance, respectively. Microbial metrics such as average richness and AWCD are positively associated with EC, while pH, Ca, and Mg are negatively associated with microbial diversity metrics. The grouping of samples highlights distinct clustering based on sample type, with water samples clustering along higher EC values and sediment samples more spread out based on nutrient concentrations (Na, P, N, K).

The first two components (PC1 and PC2) explain a significant proportion of the data variation, representing 34.6% and 19.1%, respectively. Together they account for more than 53% of the variability in the dataset. The third component (PC3) gives an additional 10.9%, resulting in a cumulative explained variance of 64.6%. Subsequent components contribute smaller amounts of variance, diminishing beyond PC4. Water samples are strongly associated with higher microbial activity and substrate utilization diversity metrics (AWCD, richness, and Shannon diversity), driven primarily by EC. In contrast, sediment and moonmilk exhibited weak associations with all variables. An inverse correlation of Na and K with richness and AWCD suggests that increased concentrations of these ions may inhibit microbial activity and substrate utilization diversity.

### Generalized Additive Models (GAMs)

The GAM plots (Figures 9, 10 and 11) revealed significant non-linear relationships between environmental variables and microbial metrics. Electrical conductivity was one of the parameters that could explain variations in microbial substrate consumption diversity. The highest Shannon diversity was observed at intermediate EC levels (100-150 µS/cm), while elevated EC levels were associated with reduced diversity. Phosphorus (P) showed a negative linear relationship with both richness and diversity, whereas sodium (Na) and magnesium (Mg) demonstrated a positive influence (Figure 9).

**Figure 9.**
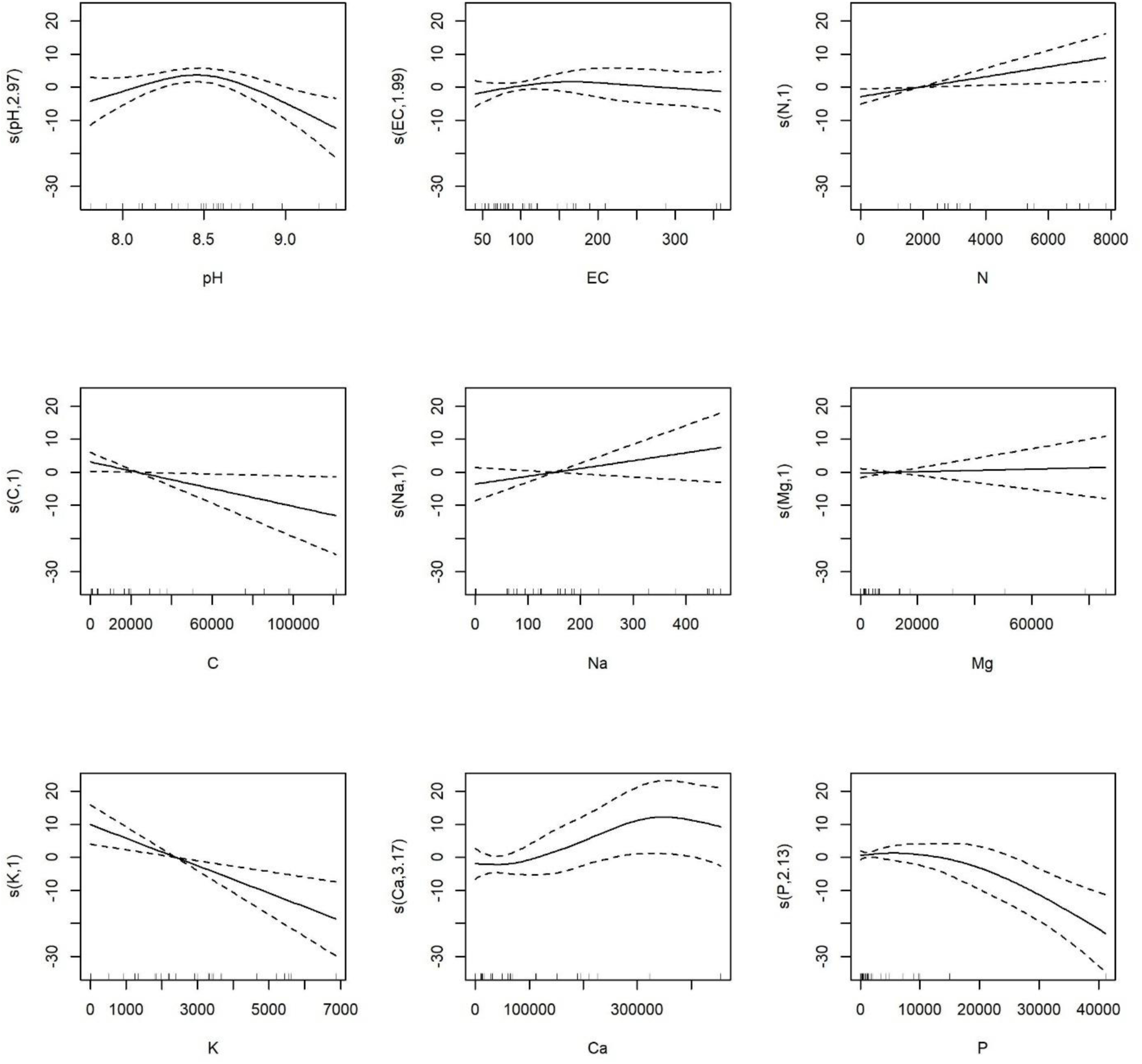
Generalized Additive Model plots showing the relationship between average richness and chemical factors (pH, EC, N, C, Na, Mg, K, Ca and P) in the analyzed cave samples. The solid lines represent the fitted smooths for each variable, while the dashed lines indicate the 95% confidence intervals. Significant relationships are observed for several variables, including a negative association between average richness, carbon and potassium, as well as non-linear effects of calcium and phosphorus on microbial substrate utilization richness.

**Figure 10.**
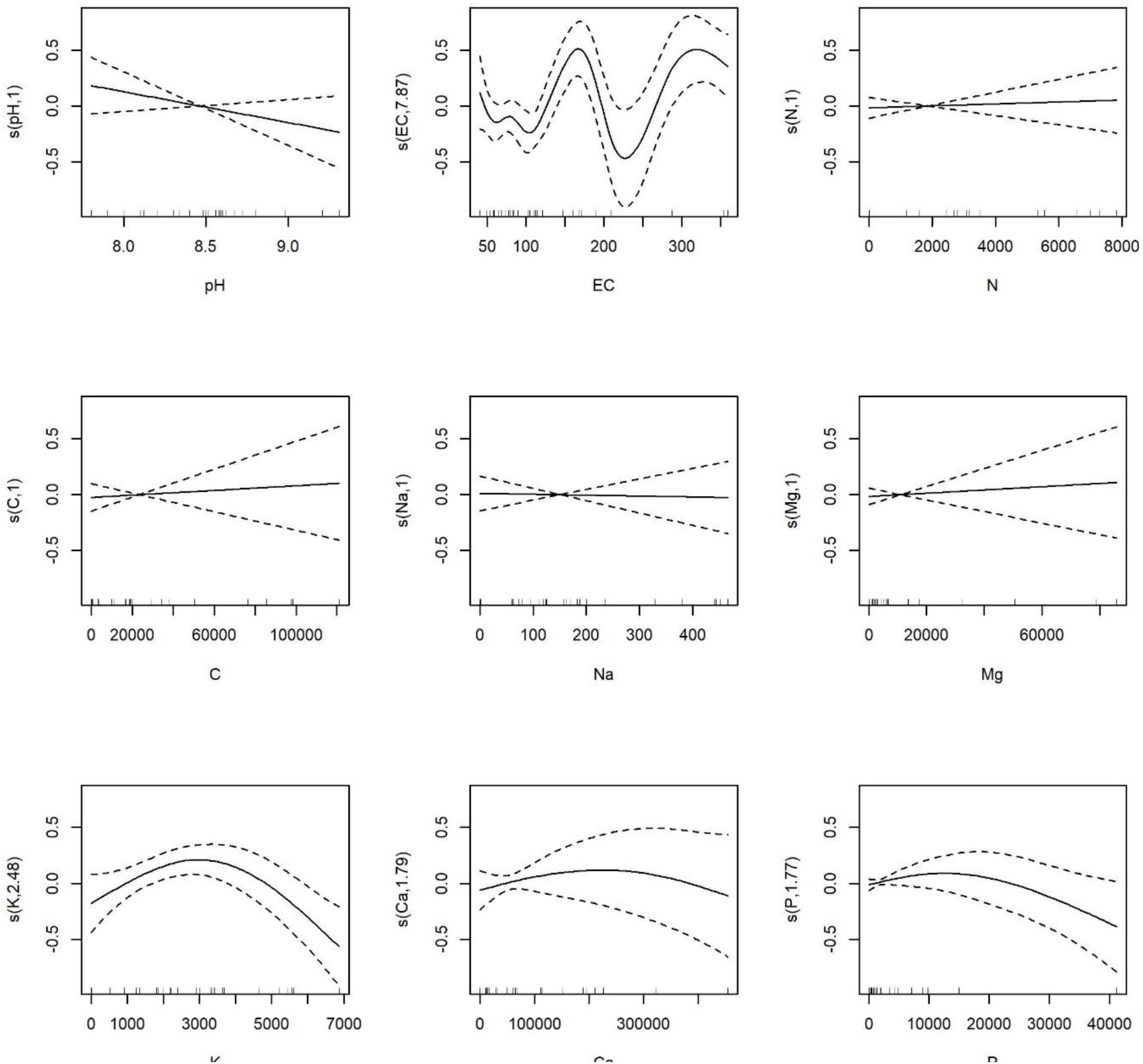
GAM plots showing the links between Shannon diversity and chemical factors. The solid lines represent the fitted smooths, and the dashed lines indicate the 95% confidence intervals. A complex, non-linear relationship is observed between Shannon diversity and EC, with multiple peaks and troughs, suggesting varying effects of ion concentrations on diversity. There are also weak associations with pH, C, and Ca, but no clear significant trends for the other variables.

**Figure 11.**
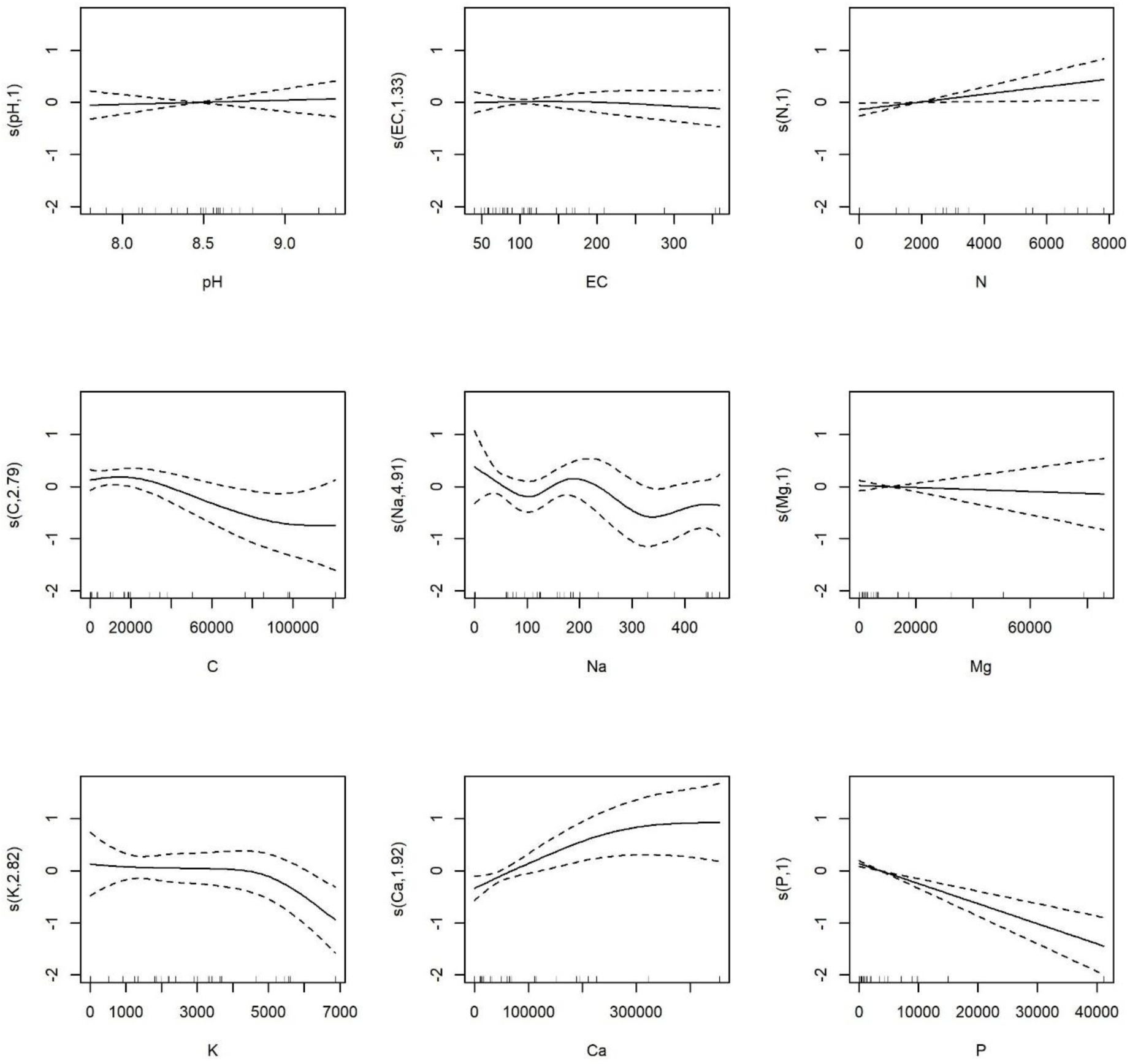
GAM plots of average Shannon evenness and different chemical factors. Solid lines – fitted smooths, dashed lines – 95% confidence intervals.

The GAM models revealed that richness and Shannon diversity are strongly influenced by chemical variables, explaining 89.1% (first model) and 91.2% (second model) of the deviance. Richness showed nonlinear relationships with pH, K, and P (p < 0.05), while N and C had significant linear effects. Shannon diversity was significantly influenced by the nonlinear effects of EC and K (p < 0.005), with P showing a trend toward significance (p = 0.077). Other variables, such as Na, Mg, and Ca, did not significantly impact richness or Shannon diversity. The correlation between Shannon diversity and EC (Figure 10) demonstrated a distinct non-linear trend, characterized by an increase in diversity at intermediate EC levels, while exhibiting a decline at both lower and higher extremes. This suggests that moderate levels of EC (likely below 100) may enhance the diversity of utilized substrates, whereas extreme levels (likely between 120-200) the response fluctuates significantly which could adversely affect the diversity. A minor negative correlation was noticed between pH and Shannon diversity, suggesting that more alkaline environments may results in a short decrease in consumed substrates diversity. Other variables, such as C, Na and Ca displayed moderate correlations with Shannon diversity, indicating their potential minimal impact on microbial community diversity in these cave samples. The final model examines the impact of chemical variables on Shannon evenness (Figure 11). The adjusted R² is 0.87, with 94.4% of the deviance explained. Key takeaways include that Shannon evenness displays a non-linear relationship with Na, suggesting that concentrations around 100 mg/kg may support a more even distribution of microbial communities, while higher (>200 mg/kg) or lower (<50 mg/kg) levels may disrupt evenness. Besides sodium (Na), potassium (K) and carbon (C) both exhibited a slight declining trend in their relationship with Shannon evenness, suggesting that higher concentrations of these elements may negatively affect evenness (Figure 11). In contrast, a positive correlation was represented by Ca, suggesting that elevated Ca levels might promote more balanced microbial communities. EC, pH and nitrogen do not significantly affect evenness.

## Discussion

### Variability and constraints in carbon substrates utilization analysis

After 200 hours of incubation, the C-substrate degradation patterns differed across water, sediment and moonmilk samples from Cloșani, Ferice, Muierilor, Leșu and Topolnița caves. Although the same volume of inoculum was applied to each well, we must take into consideration that there might be a variability in microbial inoculum size which may introduce inconsistencies in the results. Variability in microbial inoculum size likely contributed to discrepancies in optical density (OD) values, as showed by variations in wells containing the same substrate (Supplementary Figure S1). Water samples showed greater C-substrate utilization potential and thus in microbial activity compared to sediment and moonmilk samples, supporting previous studies indicating that aquatic environments rich in organic matter support metabolically diverse communities (Obusan, 2023; Tobias-Hünefeldt et al., 2023). Aquatic microbial communities are shaped by dynamic factors, including temperature and availability of organic carbon, which enhance their metabolic capabilities (Li et al., 2021; Zheng et al., 2014), suggesting that water environments can support a broader range of microbial taxa that can utilize diverse carbon sources. Thus, we could reasonably speculate on the relevance of aquatic microbial communities as key contributors to cave organic carbon degradation. Conversely, the stability of sediment and moonmilk geochemistry may restrain microbial diversity and its derived activity, with moonmilk samples displaying certain organic substrate degrading versatility. To the best of our knowledge, the present study is the first to evaluate the organic substrate utilization in moonmilk based on Biolog®EcoPlate™ approach and the few investigations on the metabolic potential of microorganisms associated with moonmilk formations prevent us from further assumptions.

The measurement of optical density in the Biolog®EcoPlate™ involves repeated handling and exposure to external environments, which increases the risk of contamination, especially for the plates incubated at temperatures higher than the room temperature (e.g., 37°C). Incubation was performed at 16°C to mitigate contamination risks and mimic cave conditions. Lower temperatures maintain microbial community structure while potentially decreasing metabolic activity, thereby improving the reliability of experimental outcomes (Adekanmbi et al., 2022; Tang et al., 2017; Akbari and Ghoshal, 2015).

### Diversity analysis based on organic substrate metabolization

Despite the individual sample variation there are more metabolically active communities associated with higher microbial metabolic richness (Figure 5). Substrate utilization patterns varied by sample type, indicating variations in nutrient availability and environmental conditions. Sediment and water samples demonstrated greater microbial community-based substrate utilization diversity, whereas moonmilk associated microbial communities displayed intermediate richness. The findings correspond with research that associates substrate richness with improved metabolic functionality across various environmental conditions (Zhang et al., 2020; Patsch et al., 2018; Li et al., 2017).

Hierarchical clustering and temporal analysis of substrate utilization showed distinct differentiation among sample types with distinct patterns across water, sediment, and moonmilk samples. Sediment samples demonstrated significant microbial metabolic versatility, employing various carbon sources as a result of the community heterogeneous composition (Meyer et al., 2022). Here, sediment samples showed moderate C-substrate utilization with clear preferences for L-serine and L-arginine, but compared to water samples indicated a much lower metabolic efficiency. Koner et al. (2021) evaluated carbon substrate use patterns in limestone caves, demonstrating that microbial communities in sediment samples exhibited diverse metabolic activity. Their findings indicate that nutrient availability affects microbial growth, with sediment samples demonstrating greater richness and diversity than water samples, which contained more specialized communities. The clustering pattern of water samples exhibited selective carbon utilization, likely influenced by nutrient-rich yet special conditions (Tobias-Hünefeldt et al., 2021; Power et al., 2018; Pašić et al., 2010). The microbial communities from moonmilk samples exhibited distinct substrate utilization patterns, especially for amino acids and amides, probably influenced by geological conditions (Nyyssönen et al., 2014).

The broad range of metabolic activity inferred to Cloșani and Muierilor caves sediments underscores their microbial diversity and metabolic potential, aligning with findings from aquifer microbiome research (Wu et al., 2015). Again, the separation of moonmilk samples in the dendrogram suggests that their distinct chemistry and environmental conditions support a specialized microbial community. Zheng and Gong (2019) and Theodorescu et al. (2023) evaluated niche differentiation within microbial communities and discovered that varying environmental conditions can result in differences in microbial diversity and composition. Their findings support the idea that microbial communities within the moonmilk samples may have distinct metabolic requirements shaped by their environmental settings.

Moreover, it is important to consider that the findings of this study provide valuable insight into the functional potential of microbial communities but represent a temporal snapshot from a single season, spring. Despite the relative temperature of the caves it might be of interest to capture seasonal variations in microbial metabolic dynamics. Additionally, the set of carbon sources of Biolog^®^ EcoPlates™ may not fully encompass the range of substrates utilized by the microbial communities, limiting insights into specific metabolic pathways or microbial identities. Future research combining metagenomics and broader substrate profiling could offer a more comprehensive understanding of the ecological roles of these communities.

### Influence of environmental chemistry on C-substrate utilization capability

The Principal Component Analysis (PCA) illustrates clustering based on sample type, associating water samples with microbial metabolic diversity metrics (Shannon diversity, evenness, richness) and electrical conductivity, whereas sediment samples showed correlation with nutrient gradients, including potassium (K), phosphorus (P), and nitrogen (N). Moonmilk samples exhibited decreased associations, indicating unique microbial responses. The findings correspond with research emphasizing the impact of dissolved ions and nutrient gradients on microbial community structure and metabolic activity (Park et al., 2020; Haegeman et al., 2013; Shaw et al., 2008). Nutrient availability facilitated microbial growth in sediments, whereas elevated sodium concentrations were inhibitory, aligning with findings from cave and reservoir systems (D’Angeli et al., 2019; Mandal et al., 2017). Electrical conductivity influenced microbial metrics in water, influencing both diversity and activity (Shen et al., 2022).

Generalized Additive Models (GAMs) further illustrated both linear (e.g., N, Na) and non-linear (e.g., K, P, Ca) influences of chemical variables on microbial metabolic diversity indices. Additionally, pH and electrical conductivity also showed non-linear effects. This underscores the significant impact of chemical environments on richness and metabolic potential of the microbial communities, similar to the study of Liu et al. (2023) in which GAMs were employed to examine the impact of different environmental variables on communities inhabiting the macrobenthos near Xiaoqing Estuary (Laizhou Bay, China) and their findings indicated that chemical factors, including salinity, organic matter, and nutrient concentrations, significantly impacted biodiversity indices that are presumably linked to the metabolic activity, revealing both linear and non-linear relationships. Moreover, the study of the phytoplankton community structure in Lake Longhu (Jiang et al., 2023) and the study of spatial patterns of biodiversity in a large marine ecosystem (Dencker et al., 2017) revealed that environmental factors, including nutrient concentrations, had significant effects on the community variability.

## Conclusions

In this study, we assessed the microbial metabolic diversity in cave sediment, water, and moonmilk samples using the Biolog® EcoPlates™ method, which enables rapid screening of carbon substrate utilization. The average-well color development (AWCD) parameter was used to evaluate substrate metabolism, while conventional diversity indices were applied to provide insights into the metabolic diversity of cave microbial communities. Hierarchical clustering revealed significant differences between sample groups, with distinct functional profiles based on the sample’s origin and geochemistry. Notably, this study is the first to apply Biolog® EcoPlates™ to moonmilk samples from karstic caves, highlighting the significance of employing novel approaches to explore the metabolic potential of microbial communities in cave environments.

The results revealed that microbial communities populating the cave water samples were the most metabolically versatile, utilizing a broad range of carbon substrates compared to those in sediment and moonmilk samples. D-galacturonic acid, L-asparagine, and Tween 80 were the most readily degraded C-substrates. The AWCD and Shannon diversity indices further suggested that cave water hosted the most metabolically diverse microbial communities, playing a critical role in carbon turnover in studied cave systems. Sediment-associated communities exhibited more restricted metabolic profiles, favoring specific substrates like L-serine and L-arginine. In contrast, moonmilk samples showed the least metabolic activity, with a narrow substrate range, indicating a specialized microbial community. Additionally, the application of Generalized Additive Models (GAMs) in cave microbiology is novel, providing a powerful approach for analyzing complex relationships between microbial metabolic diversity and environmental factors. GAMs revealed a positive correlation between Shannon diversity and intermediate values of electrical conductivity, indicating that moderate EC levels enhance microbial C-utilization.

Overall, this work offers fresh insights into the metabolic capacities of cave microbial communities and their adaptation to environmental variables, shedding light on their ecological significance in carbon cycling in cave environments.

## Acknowledgements and funding

This work was supported by a grant of the Ministry of Research, Innovation and Digitization, CNCS/CCCDI – UEFISCDI, project number 2/2019 (DARKFOOD), within PNCDI III. DFP has received financial support through the project: Entrepreneurship for innovation through doctoral and postdoctoral research, POCU/380/6/13/123886 co-financed by the European Social Fund, through the Operational Program for Human Capital 2014-2020.

We thank to Ruxandra Bucur, Cristian Sitar, Alexandru Petculescu, Ionuț Cornel Mirea and Răzvan Arghir for their contribution to sampling campaigns.

## Author contributions

– DFP, HLB, OTM designed the research and drafted the manuscript.
– DFP and AC conducted the research.
– AMP performed the statistical analyses.
– OTM performed the sampling
– EAL performed the chemical analyses

All authors contributed, verified, and approved the contents of the manuscript.

## Supplementary materials

**Figure S1.**
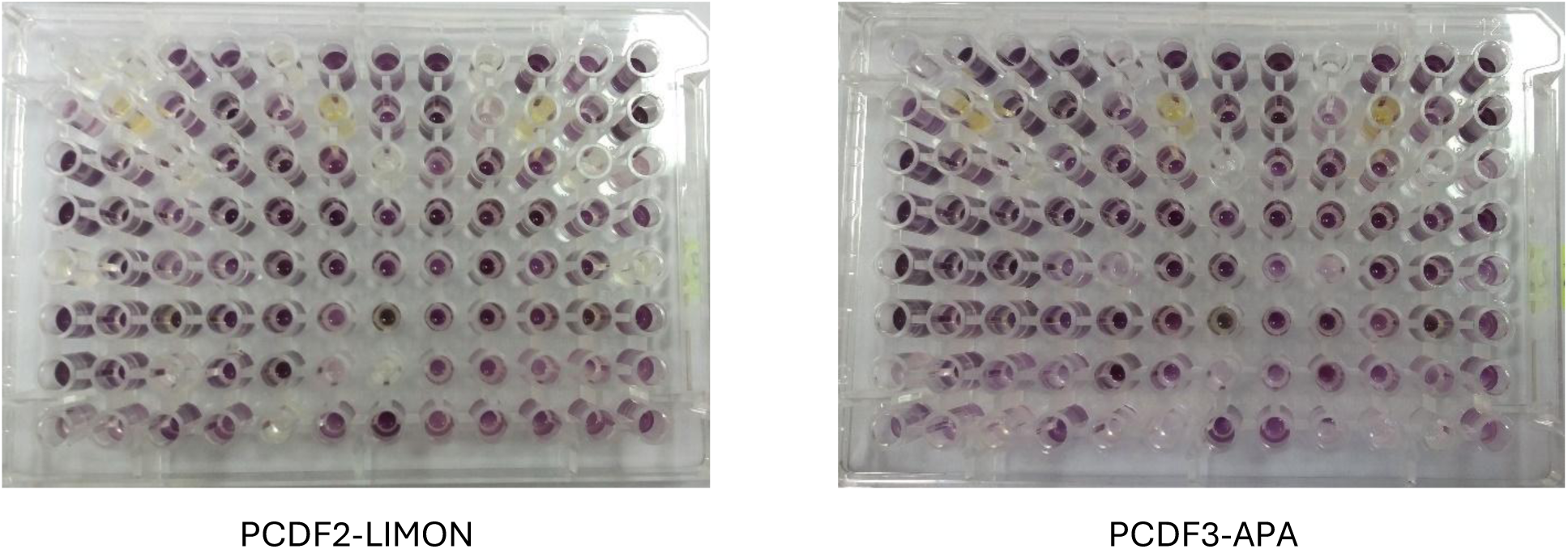
Random selection of pictures showing the degradation patterns within Biolog® EcoPlates™ after an incubation period of 200 hours (PC – Closani Cave)

**Figure S2.**
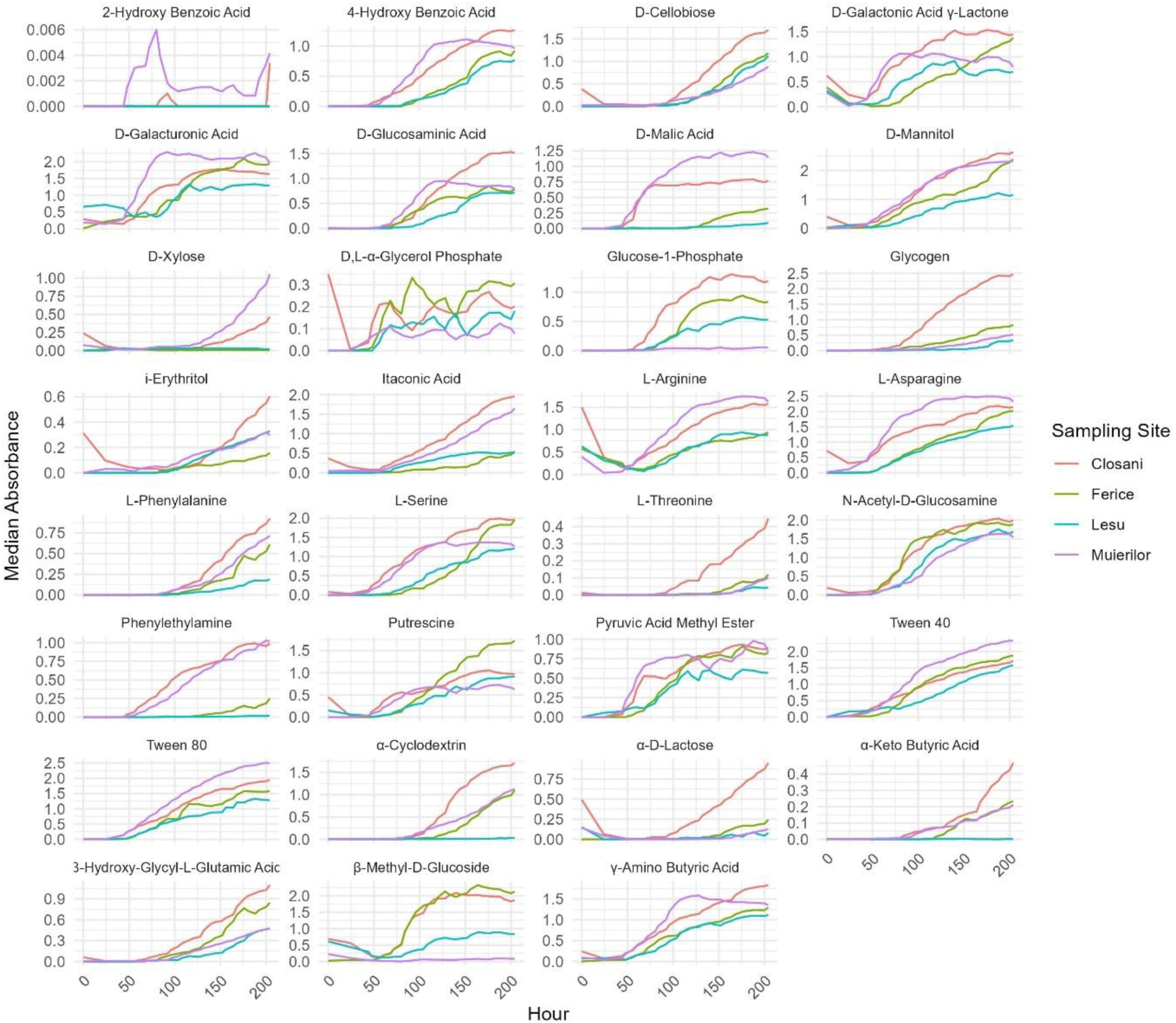
Median absorbance over time by carbon and sampling site for water samples only.

**Figure S3.**
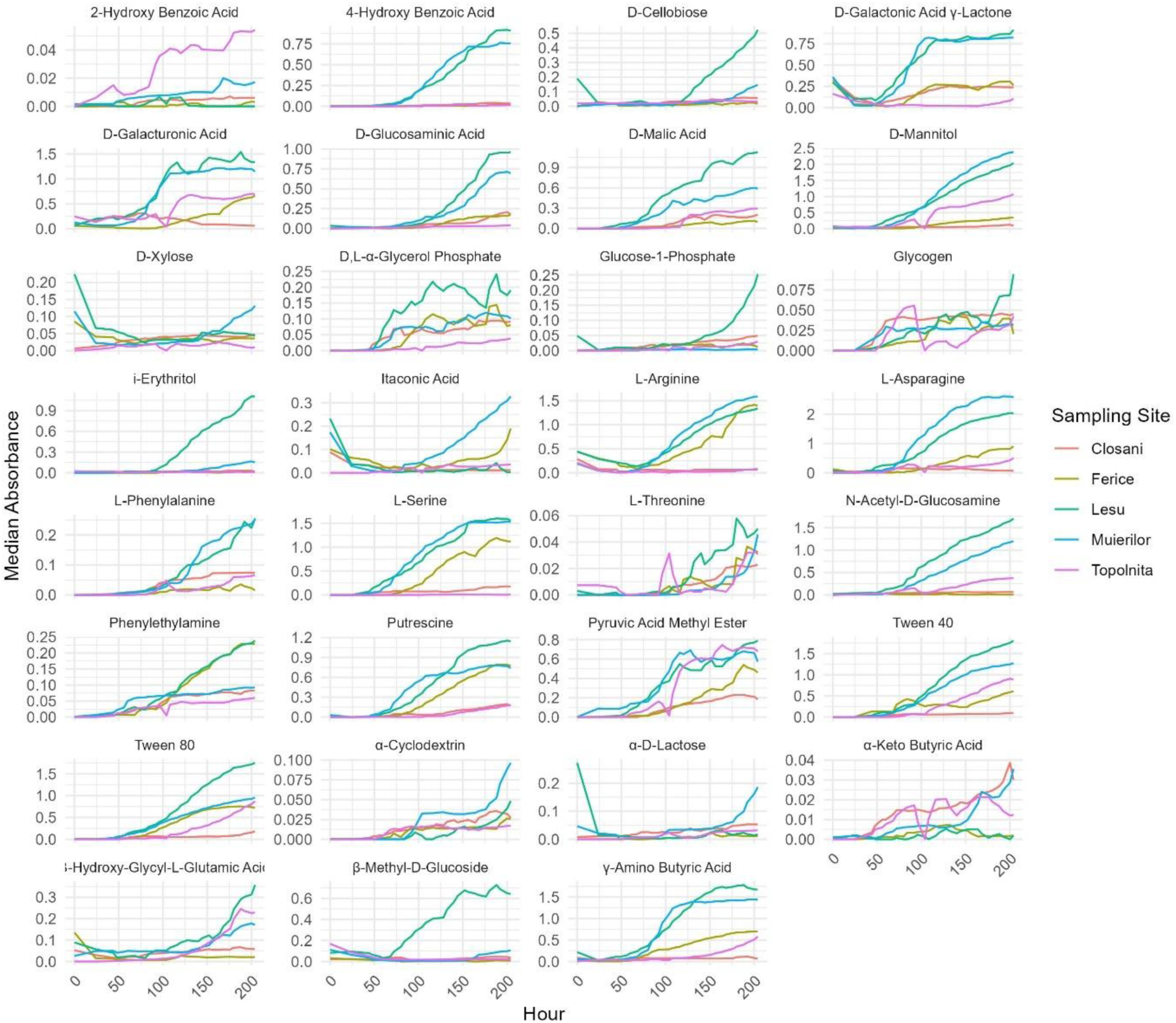
Median absorbance over time by carbon source and sampling site – sediment samples only.

**Figure S4.**
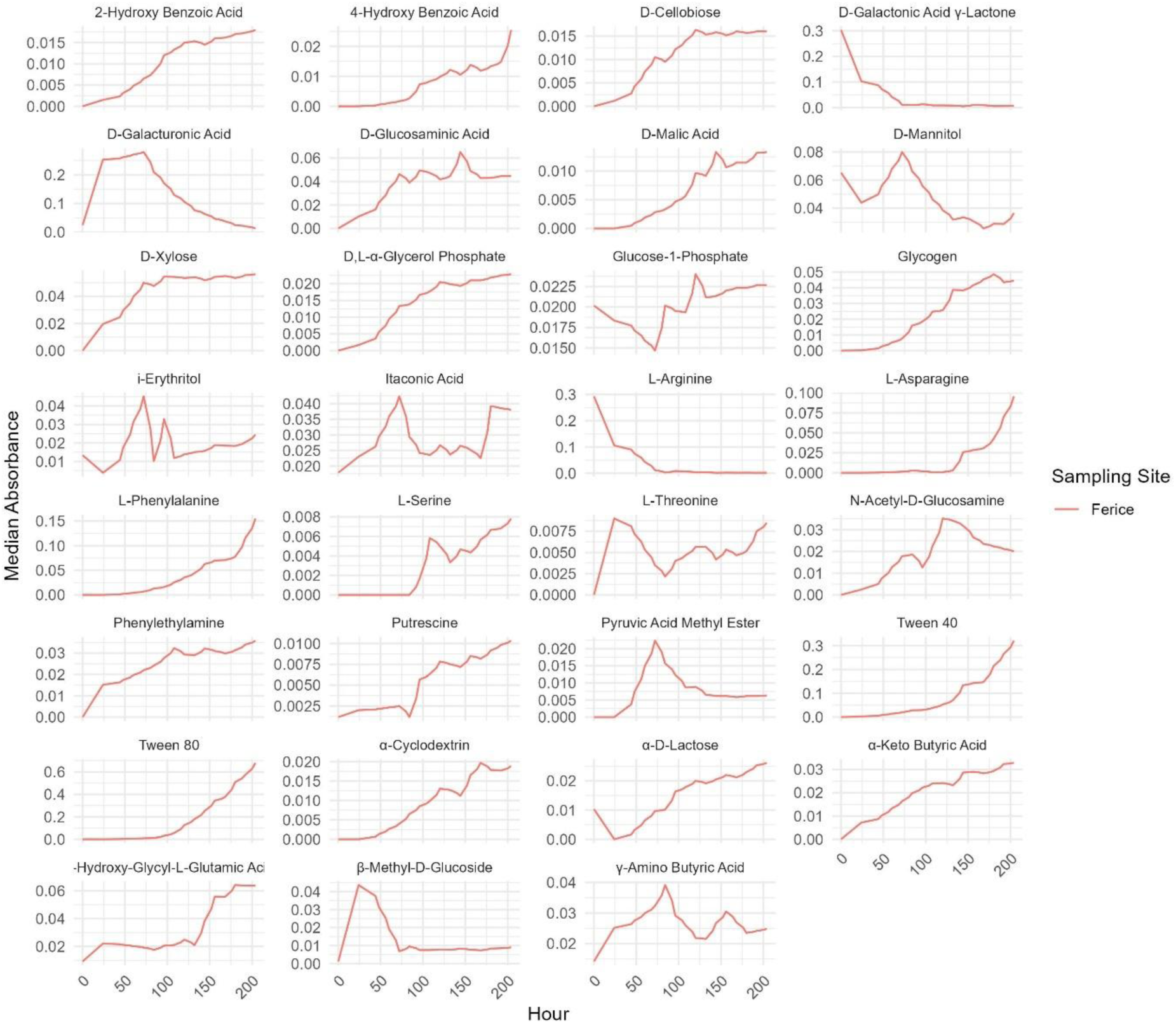
Median absorbance over time by carbon source and sampling site – moonmilk samples only.

**Figure S5.**
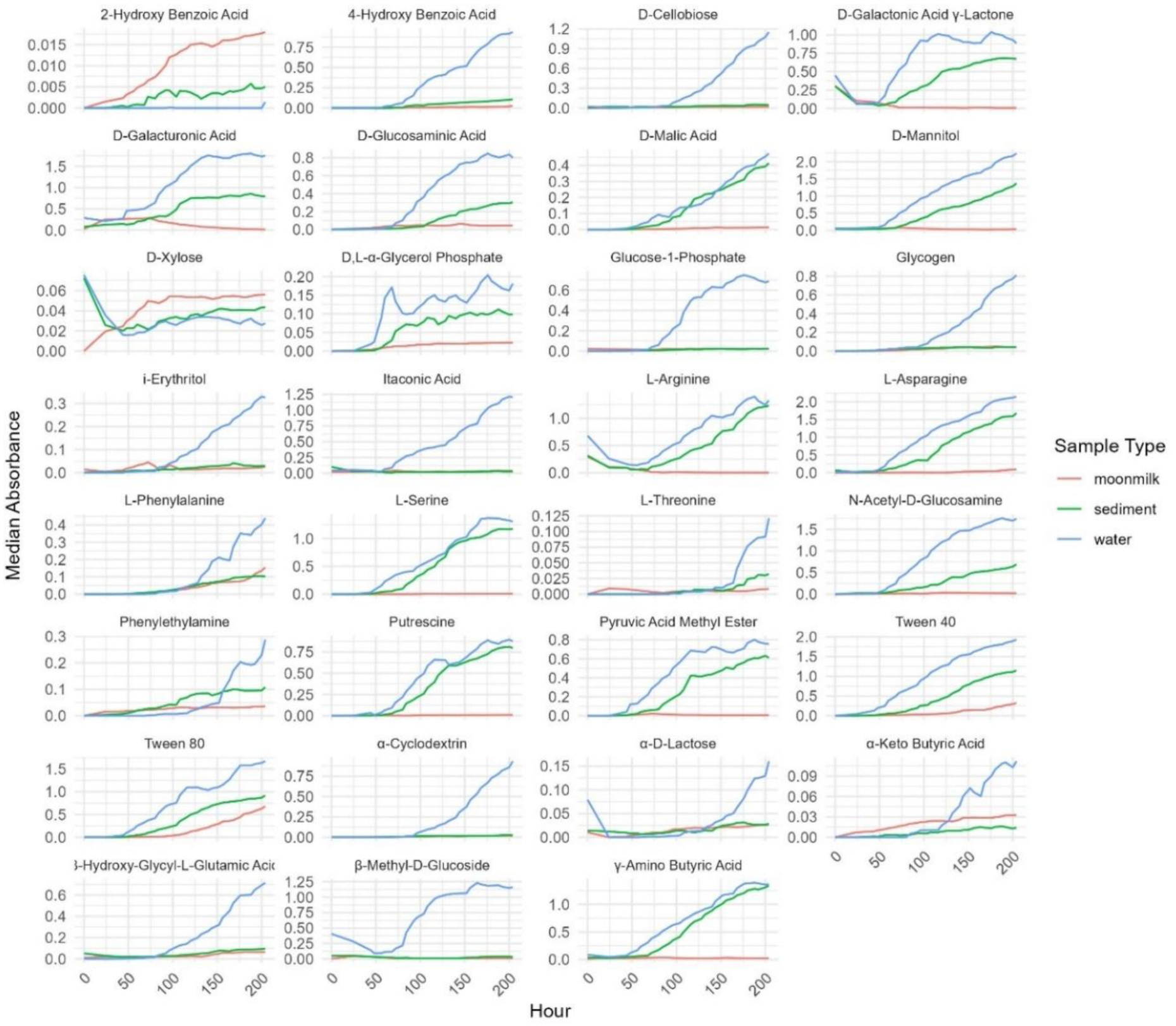
Line plots illustrate the median absorbance over time for 31 carbon substrates categorized into three sample types: moonmilk (red), sediment (green), and water (blue). Each panel corresponds to a specific carbon source, illustrating variations in the metabolic activity dynamics among the sample types.

**Figure S6.**
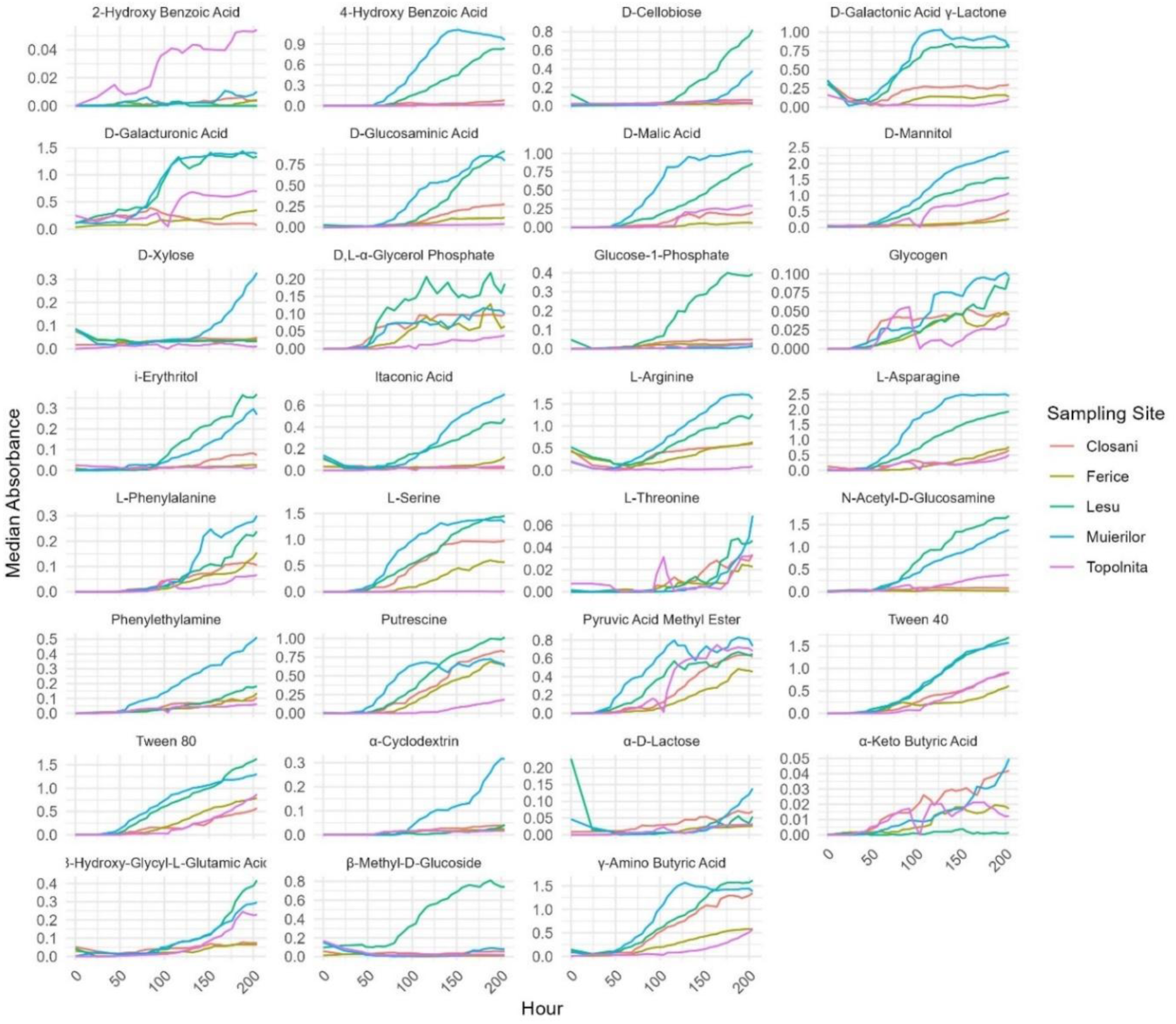
Line plots showing the median absorbance over time for the 31 C-sources among microbial communities from five sample locations: Cloșani, Ferice, Leșu, Muierilor and Topolnița Cave. Each panel represents a distinct carbon source, emphasizing site-specific variations in metabolic processes within the microbial communities.

**Table S1.** Maximum AWCD values for water, sediment, and moonmilk samples across the sampling sites.

| Site | Max AWCD<br>(Water) | Max AWCD<br>(Sediment) | Max AWCD<br>(Moonmilk) |
| --- | --- | --- | --- |
| Cloșani | 1.67 | 1.32 | - |
| Ferice | 1.01 | 1.01 | 0.07 |
| Leșu | 0.95 | 1.07 | - |
| Muierilor | 1.08 | 0.84 | - |
| Topolnița | - | 0.41 | - |

**Figure S7.**
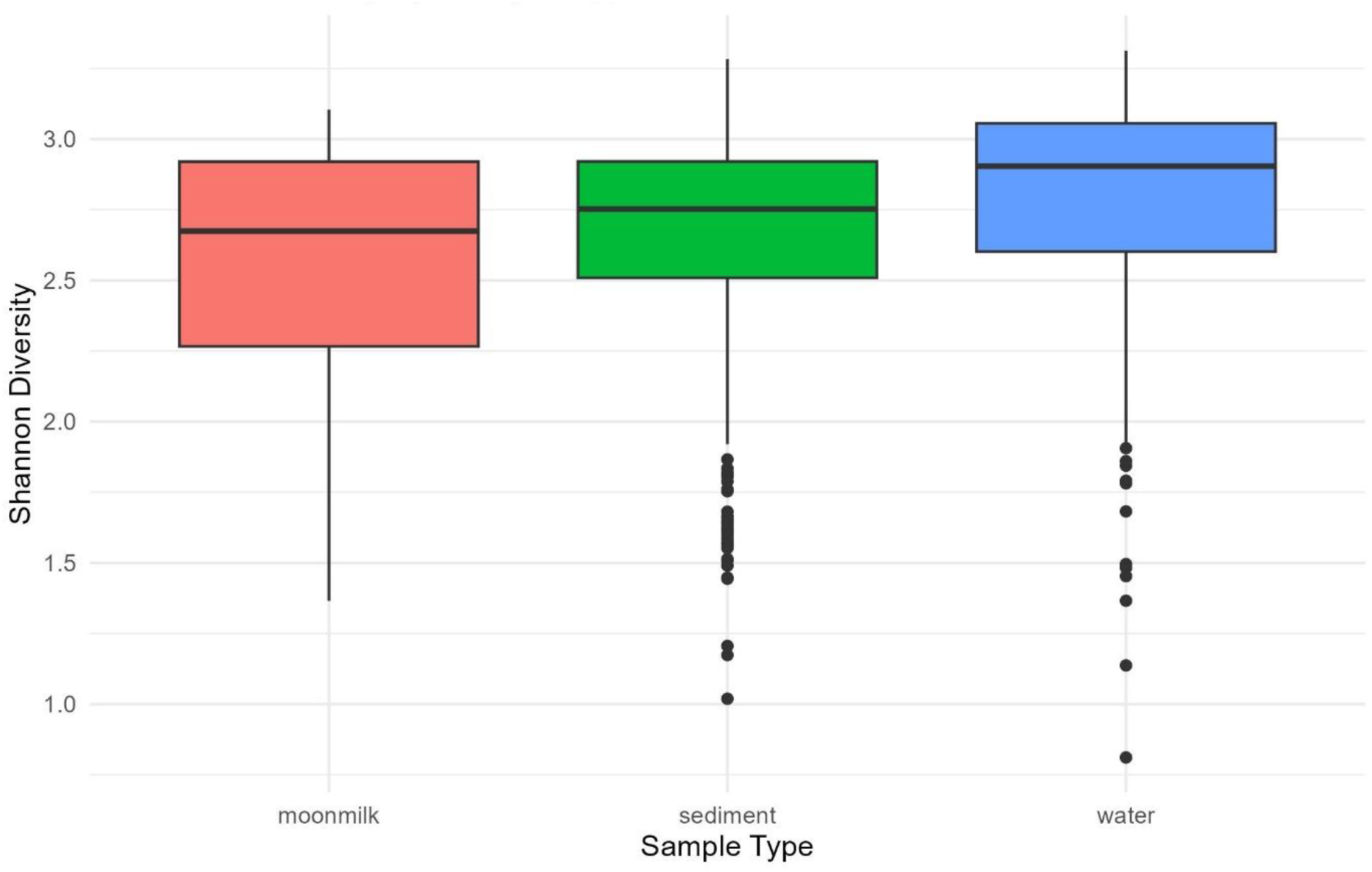
Shannon diversity (metabolic diversity) index of microbial communities in moonmilk, sediment, and water samples. Water samples show the highest diversity, followed by moonmilk and sediments. Boxplots display median, interquartile range, and outliers.

**Figure S8.**
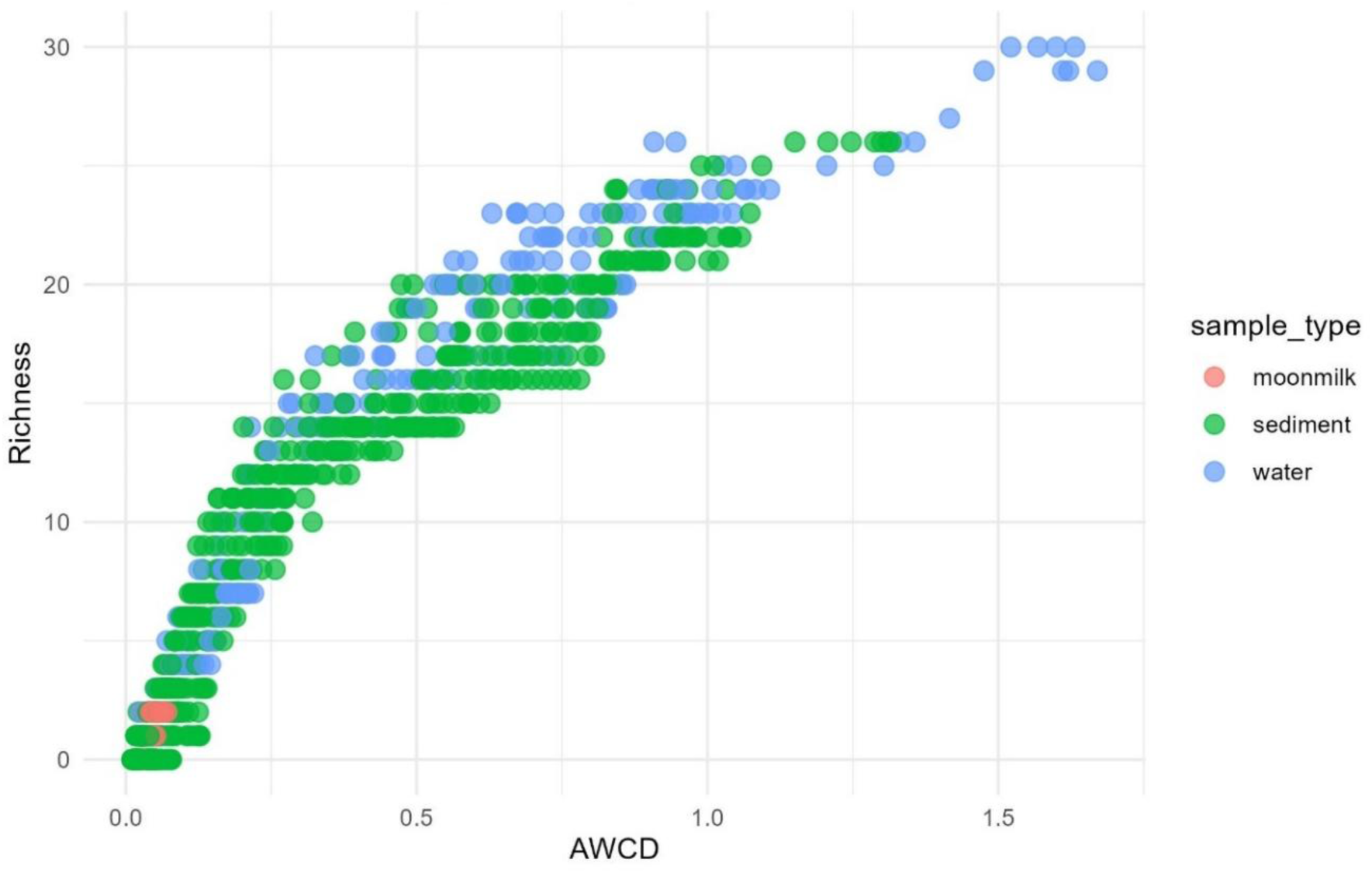
Scatter plot of AWCD versus richness of substrate degradation for sediments, moonmilk, and water samples, measured over time (0–204 hours).

**Figure S9.**
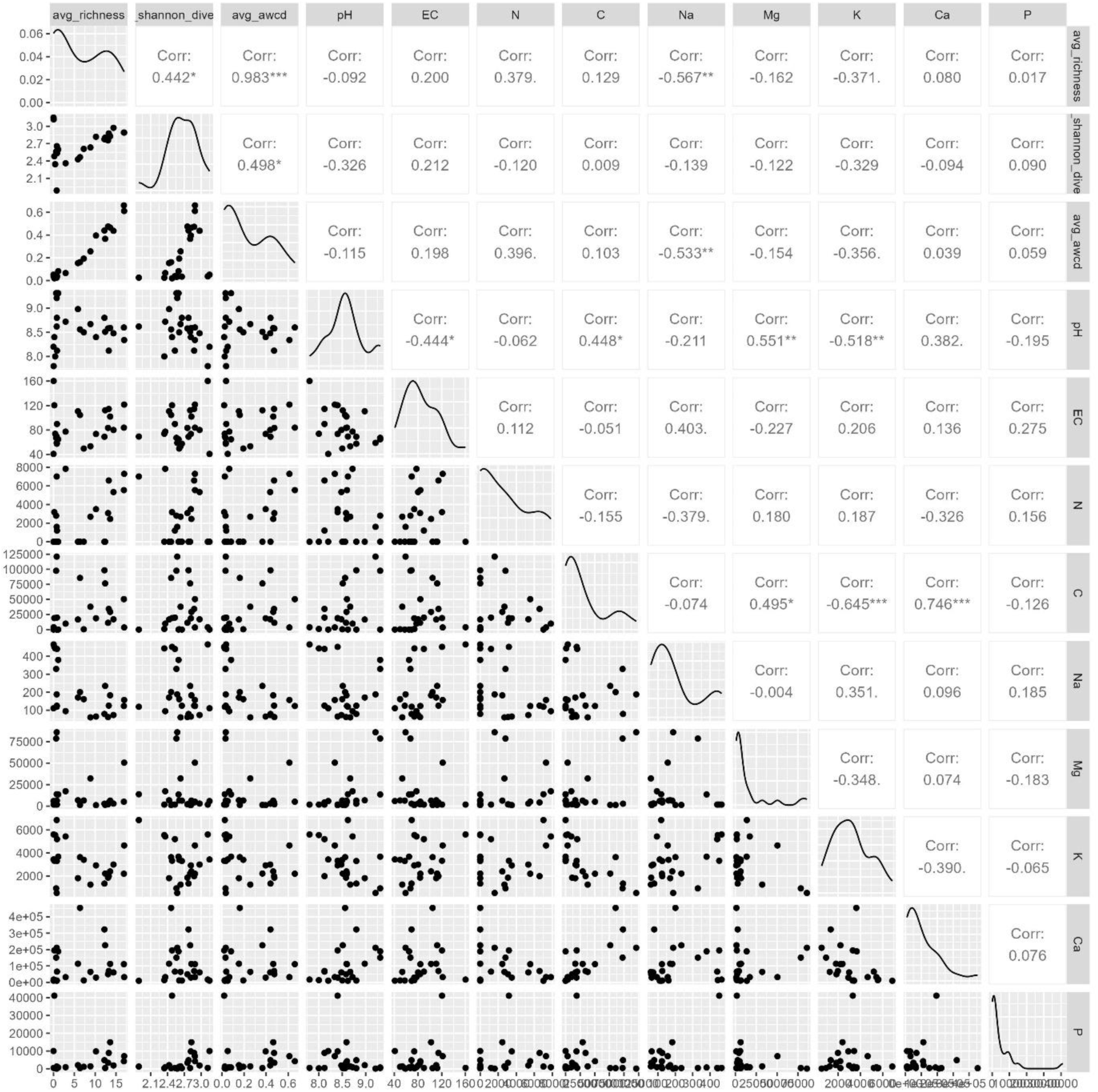
Scatterplot matrix showing the relationship between cave chemistry variables (pH, EC, nitrogen, carbon, sodium, magnesium, potassium, calcium and phosphorus) and biological metrics (average richness, Shannon diversity, average AWCD) for sediment samples. Each subplot includes a correlation coefficient (Corr), indicating the strength and direction of the relationships. Notable correlations include a positive correlation between Shannon diversity and average AWCD (r = 0.983, p < 0.001) and negative correlations between sodium (Na) and microbial metrics, such as average richness and Shannon diversity (r = –0.567 and r = –0.533, respectively). The diagonal plots represent kernel density estimates for each variable.

**Figure S10.**
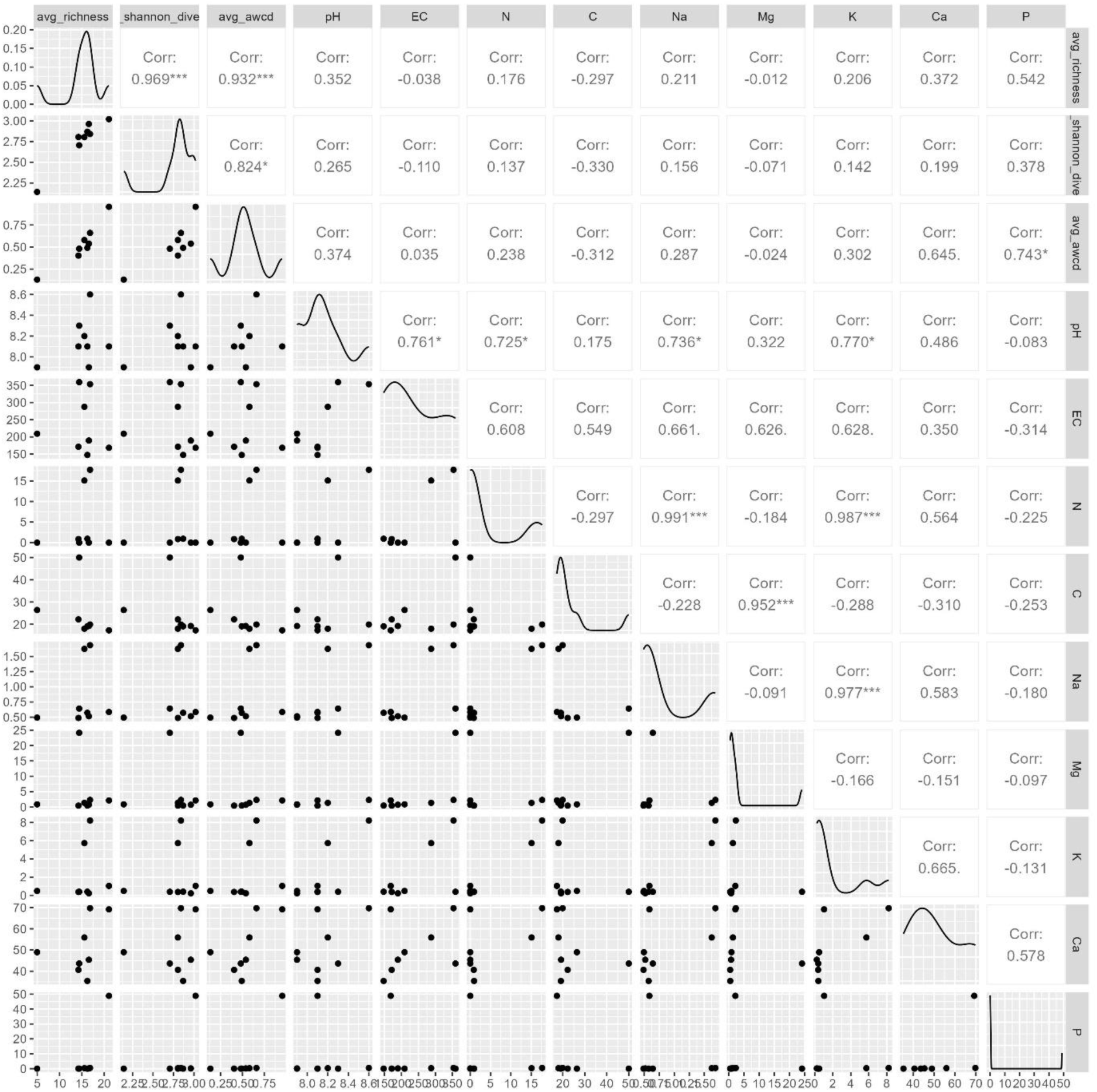
Scatterplot matrix for microbial metrics (identical as the metrics used for sediments) and cave chemistry variables (identical as the variables used for sediments) for water samples. he correlation coefficients (Corr) indicate significant positive correlations, including between Shannon diversity and average AWCD (r = 0.932, p < 0.001), as well as between electrical conductivity (EC) and various microbial metrics, such as Shannon diversity and average AWCD (r = 0.761 and r = 0.725, respectively). The kernel density plots on the diagonal represent the distribution of each variable.

**Figure S11.**
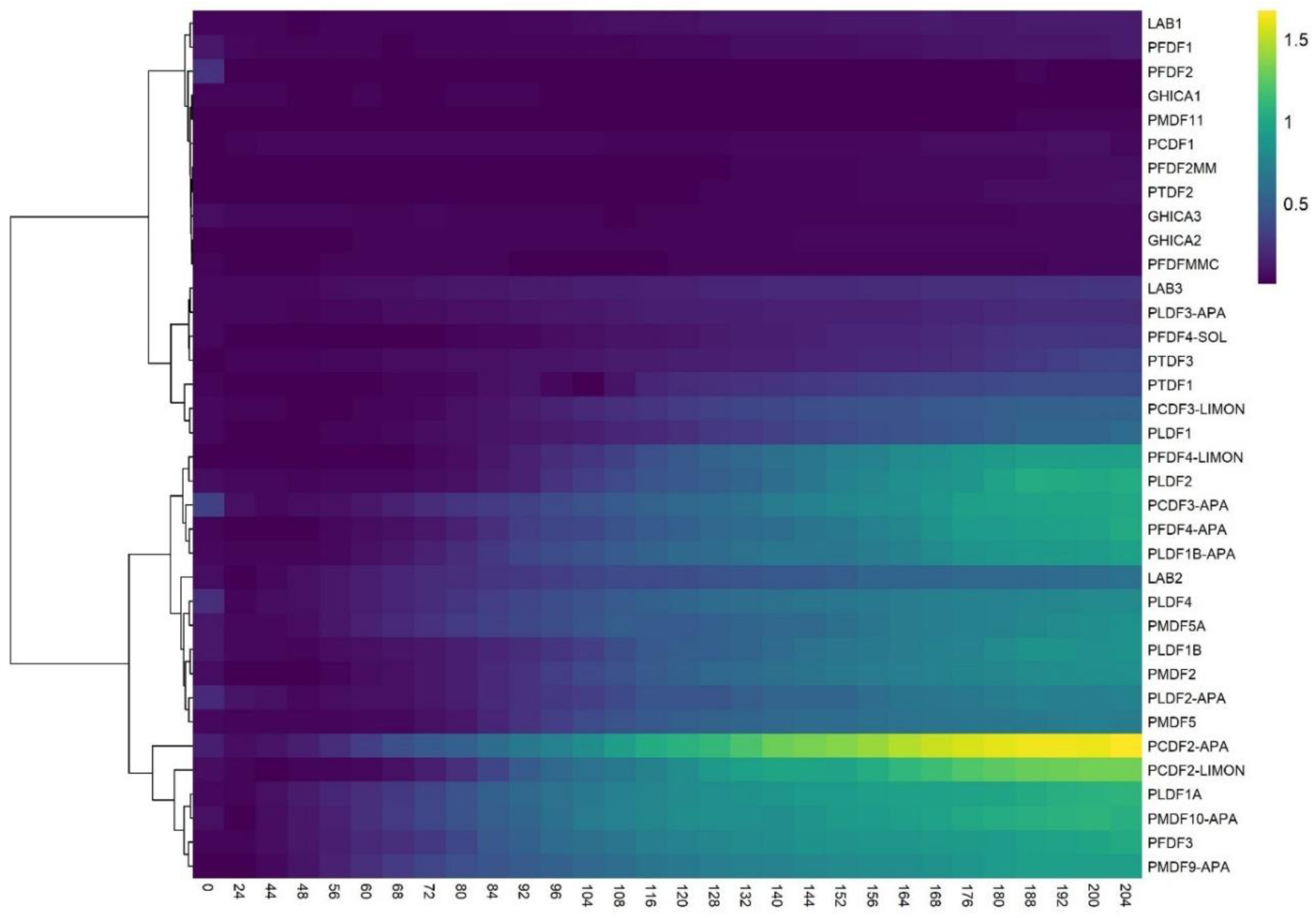
Heatmap of the average-well color development by sample and hour with clustering. See Table 1 for corresponding sample types and codes.

**Figure S12.**
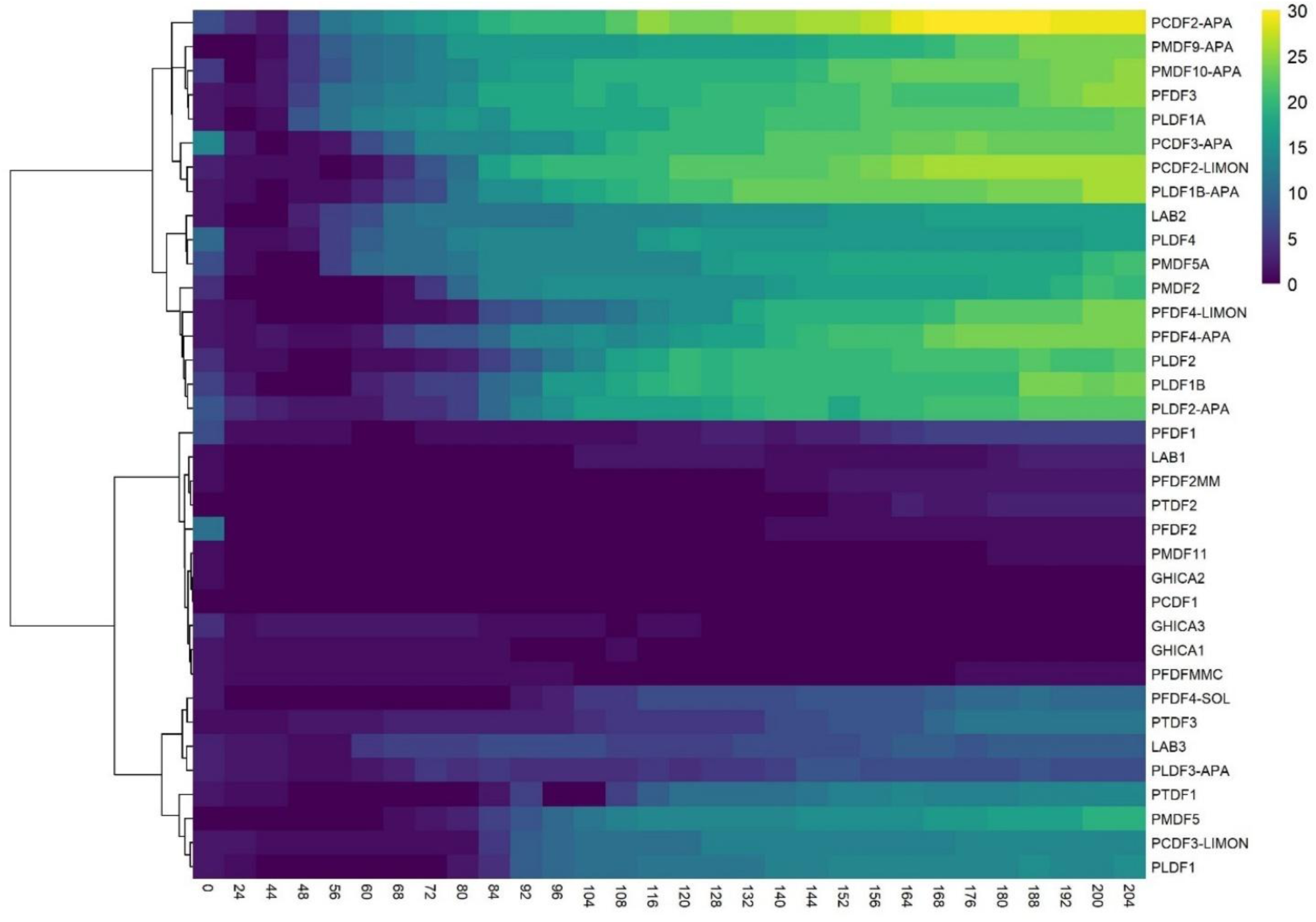
The heatmap showing the clustering of substrate richness by sample and hour. All the sample types are included in the analysis. The color scale represents species richness, with yellow indicating higher richness and dark purple representing lower richness. Hierarchical clustering was applied to both samples (rows) and time points (columns), revealing distinct temporal and sample-specific patterns in microbial diversity. See Table II.1 for corresponding sample types and codes.

**Figure S13.**
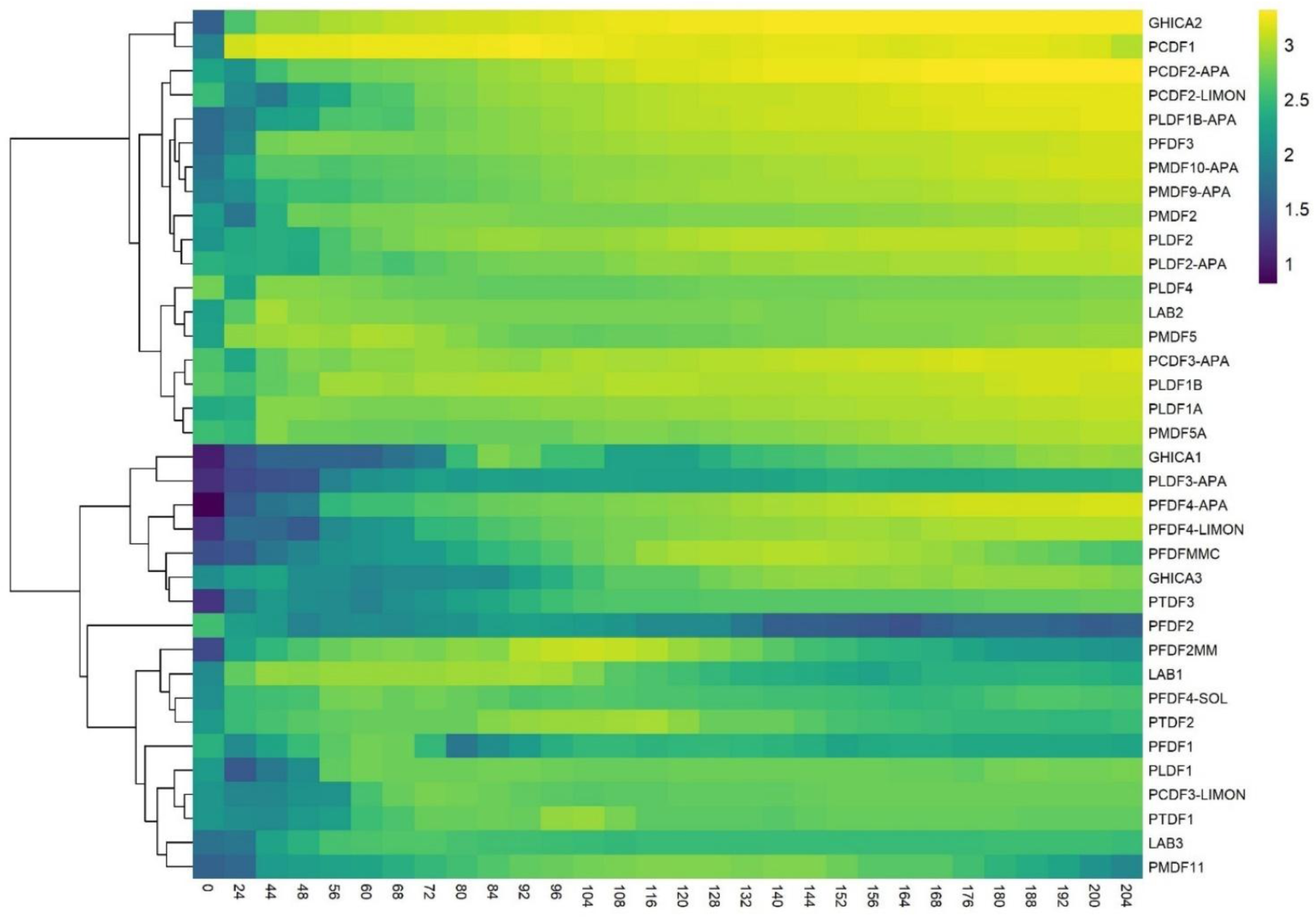
Dendrogram heatmap showing the clustering of Shannon diversity across different samples over time. The color scale represents Shannon diversity, with yellow indicating higher diversity and purple representing lower diversity. Hierarchical clustering was applied to both samples (rows) and time points (columns), revealing distinct temporal and sample-specific patterns in microbial diversity.

**Figure S14.**
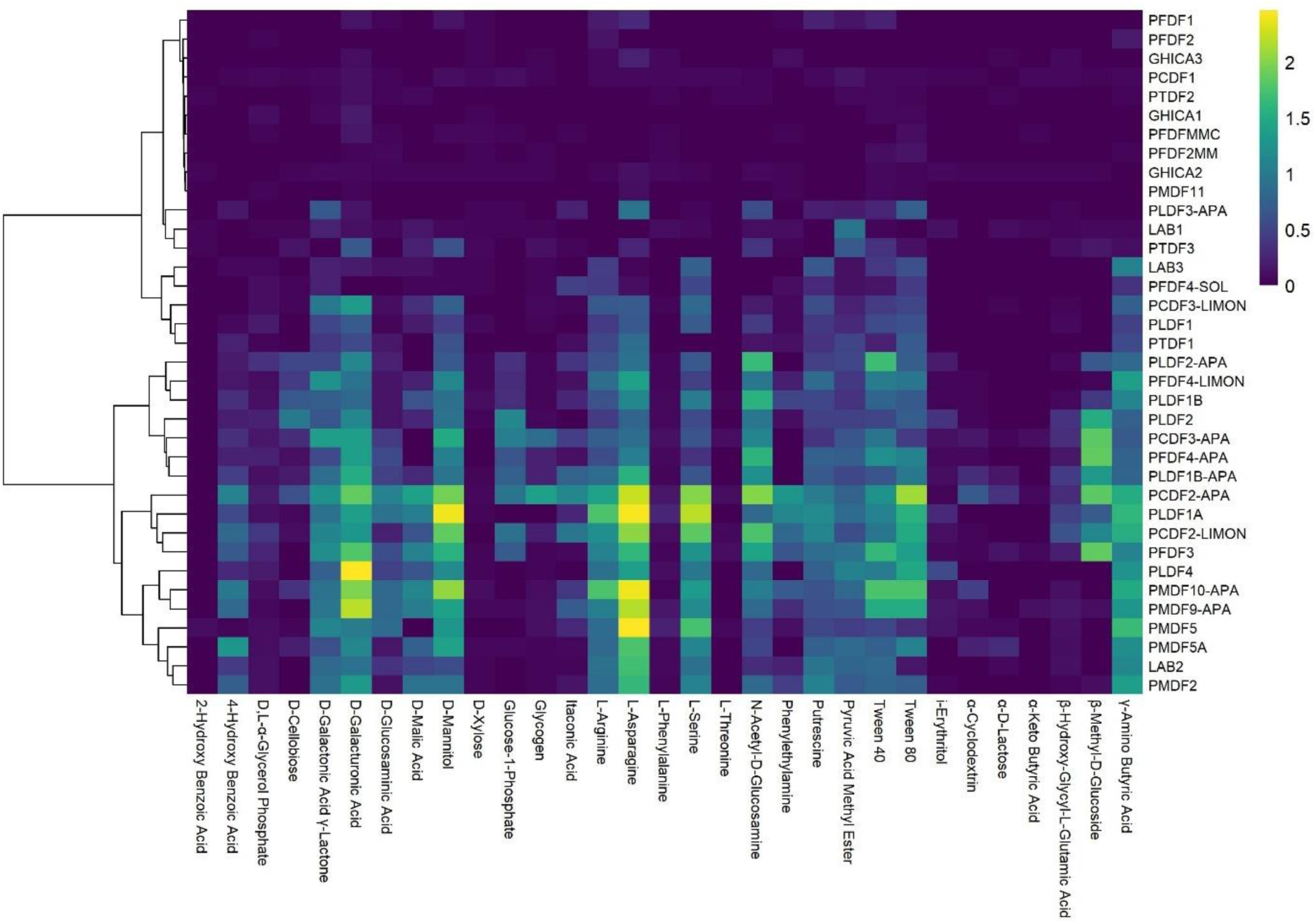
Heatmap of hierarchical clustering of samples and carbon sources based on median absorbance. Yellow indicates higher metabolic activity, purple indicates lower activity, highlighting carbon utilization patterns.

**Figure S15.**
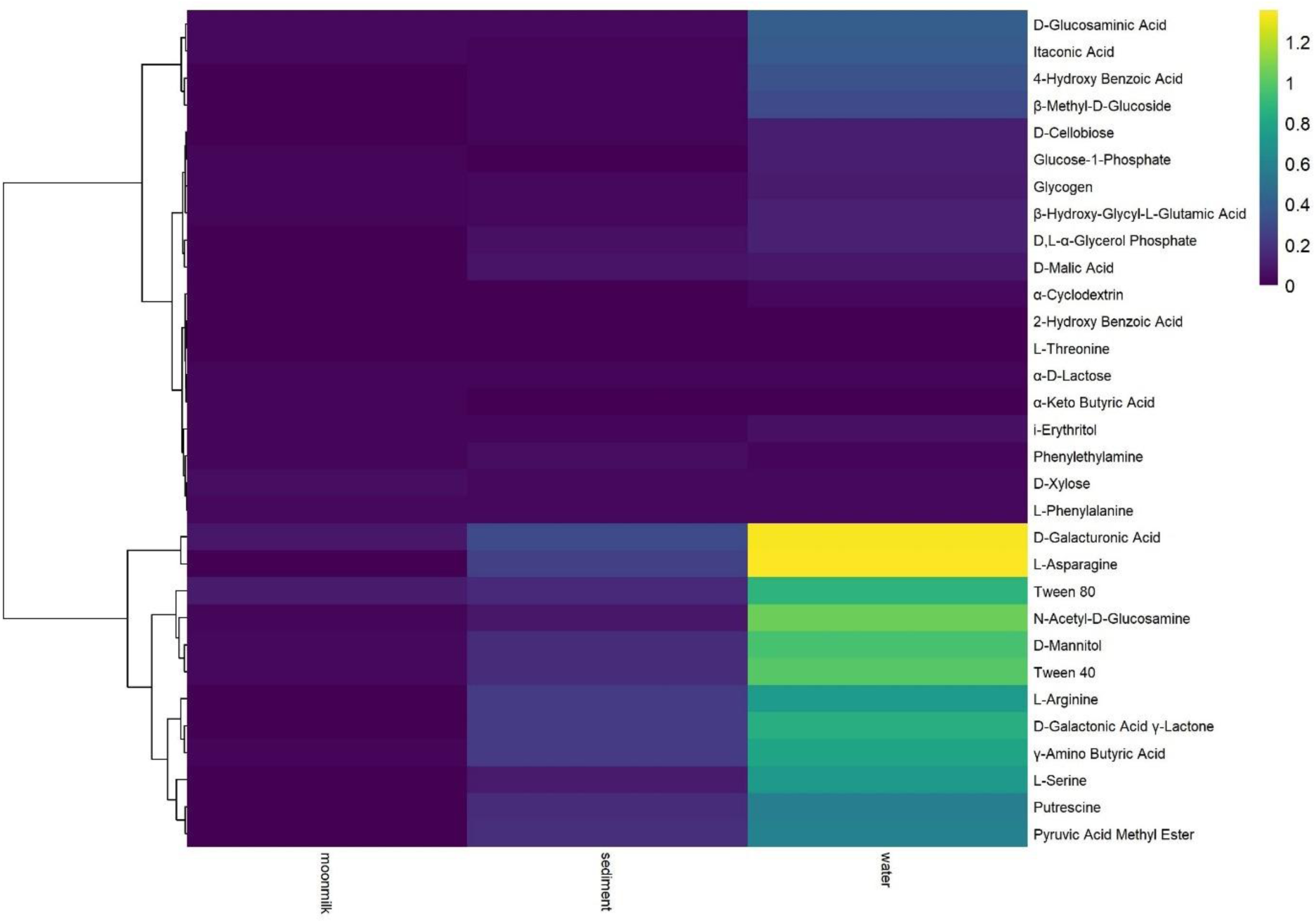
Heatmap of sample types (moonmilk, sediment, water) and carbon sources clustered by median absorbance. Yellow indicates higher and purple lower metabolic activity, showing carbon utilization patterns.

**Table S2.** List of geochemical parameters excluded from the PCA analysis, including elements (Al, Fe, S, Mn, As, Cr, Co, Ni, Cu, Zn) and water-specific parameters (DIC, DOC, SO₄^2-^, NO₃^−^, Cl^−^, PO₄^3-^), excluded for either low concentrations or incompatibility between sediment and water sample data.

| Parameters |  |  |  |  |  |  |  |  |  |  |  |  |  |  |  |  |
| --- | --- | --- | --- | --- | --- | --- | --- | --- | --- | --- | --- | --- | --- | --- | --- | --- |
| | Al | Fe | S | Mn | As | Cr | Co | Ni | Cu | Zn | DIC | DOC | $\text{SO}_4^{2-}$ | $\text{NO}_3^-$ | $\text{Cl}^-$ | $\text{PO}_4^{3-}$ |
| sediments |  |  |  |  |  |  |  |  |  |  |  |  |  |  |  |  |
| Sample ID | mg/kg | mg/kg | mg/kg | mg/kg | mg/kg | mg/kg | mg/kg | mg/kg | mg/kg | mg/kg | - | - | - | - | - | - |
| PMDF2 | 3903 | 4617 | 68.4 | 214 | 52.7 | 4.0 | 1.3 | 4.6 | 11.0 | 60.3 | - | - | - | - | - | - |
| PMDF5 | 14847 | 27613 | 196 | 158 | 15.6 | 38.0 | 3.4 | 9.3 | 467 | 242 | - | - | - | - | - | - |
| PMDF11 | 14370 | 14303 | 286 | 570 | 33.5 | 17.0 | 3.1 | 12.3 | 19.8 | 784 | - | - | - | - | - | - |
| PMDF5A | 16427 | 23567 | 322 | 195 | 8.2 | 6.0 | 1.8 | 6.4.0 | 35.1 | 56.0 | - | - | - | - | - | - |
| PTDF1 | 24903 | 33967 | 92.7 | 586 | 6.1 | 66.0 | 12.7 | 29.3 | 22.6 | 83.3 | - | - | - | - | - | - |
| PTDF2 | 25773 | 34367 | 37 | 635 | 5.5 | 59.0 | 12.9 | 28.2 | 21.0 | 64.6 | - | - | - | - | - | - |
| PTDF3 | 14866 | 23100 | 206 | 81.2 | 5.3 | 8.5 | 1.7 | 5.0 | 3.4 | 9.2 | - | - | - | - | - | - |
| LAB1 | 33362 | 30139 | 27.6 | 1097 | 5.1 | 49.7 | 19.7 | 51.7 | 51.2 | 156 | - | - | - | - | - | - |
| LAB2 | 24540 | 22837 | 46.7 | 186 | 13.5 | 42.2 | 6.7 | 40.2 | 28.9 | 489 | - | - | - | - | - | - |
| LAB3 | 38107 | 28104 | 39.6 | 138 | 11.9 | 34.0 | 5.2 | 32.5 | 21.5 | 153 | - | - | - | - | - | - |
| GHICA1 | 62997 | 55572 | 61.1 | 313 | 13.9 | 93.7 | 8.2 | 69.7 | 46.4 | 148 | - | - | - | - | - | - |
| GHICA2 | 63089 | 53824 | 85.8 | 305 | 14.1 | 89.7 | 8.2 | 72.8 | 47.1 | 154 | - | - | - | - | - | - |
| GHICA3 | 57706 | 45181 | 47.3 | 304 | 13.1 | 89.0 | 8.1 | 73.1 | 49.6 | 156 | - | - | - | - | - | - |
| PCDF1 | 27557 | 39667 | 30.9 | 945 | 6.2 | 44.0 | 14.1 | 30.0 | 32.7 | 96.5 | - | - | - | - | - | - |
| PFDF1 | 28283 | 32257 | 59.0 | 622 | 20.2 | 38.0 | 7.7 | 27.0 | 15.8 | 130 | - | - | - | - | - | - |
| PFDF2 | 25933 | 31157 | 31.8 | 998 | 17.2 | 38.0 | 10.6 | 31.5 | 17.1 | 111 | - | - | - | - | - | - |
| PFDF3 | 40033 | 32097 | 67.8 | 686 | 19.7 | 46.0 | 11.0 | 23.9 | 17.6 | 170 | - | - | - | - | - | - |
| PFDFMMC | 12768 | 6089 | 24.5 | 128 | 29.6 | 13.4 | 1.8 | 17.6 | 1.6 | 31.3 | - | - | - | - | - | - |
| PFDF2MM | 3286 | 1853 | 86.2 | 66.3 | 5.6 | 5.6 | 1.2 | 6.0 | 0.3 | 14.7 | - | - | - | - | - | - |
|  | Al | Fe | S | Mn | As | Cr | Co | Ni | Cu | Zn | DIC | DOC | SO<sub>4</sub><sup>2-</sup> | NO<sub>3</sub><sup>-</sup> | Cl<sup>-</sup> | PO<sub>4</sub><sup>3-</sup> |
| sediments |  |  |  |  |  |  |  |  |  |  |  |  |  |  |  |  |
| Sample ID | mg/kg | mg/kg | mg/kg | mg/kg | mg/kg | mg/kg | mg/kg | mg/kg | mg/kg | mg/kg | - | - | - | - | - | - |
| PLDF1 | 9543 | 16157 | 36 | 411 | 23.8 | 13 | 5.5 | 13.9 | 5.5 | 28.5 | - | - | - | - | - | - |
| PLDF1A | 19853 | 26813 | 734 | 161 | 13.4 | 5 | 1.5 | 5.0 | 5.4 | 16.3 | - | - | - | - | - | - |
| PLDF1B | 18113 | 25193 | 176 | 832 | 53.3 | 21 | 8.3 | 27.8 | 26.0 | 134 | - | - | - | - | - | - |
| PLDF4 | 12183 | 22273 | 128 | 745 | 51.0 | 19 | 7.3 | 25.1 | 28.5 | 134 | - | - | - | - | - | - |
| PLDF2 | 18173 | 27787 | 801 | 272 | 13.8 | 4.7 | 1.8 | 5.9 | 8.7 | 51.4 | - | - | - | - | - | - |
| water |  |  |  |  |  |  |  |  |  |  |  |  |  |  |  |  |
|  | µg/L | µg/L | - | µg/L | µg/L | µg/L | µg/L | µg/L | µg/L | µg/L | mg/L | mg/L | mg/L | mg/L | mg/L | mg/L |
| PMDF9-APA | <0.2 | 30.0 | NA | 0.20 | 0.50 | 1.00 | <0.2 | <0.2 | 0.7 | 5.1 | 16.9 | 3.3 | 13.5 | 61.5 | 1.9 | 13.5 |
| PMDF10-APA | <0.2 | 30.0 | NA | 0.20 | 0.80 | 1.20 | <0.2 | 0.4 | 0.9 | 3.5 | 19.8 | 2.1 | 28.0 | 78.5 | 2.7 | 28.0 |
| PCDF2-APA | 7.8 | 6800 | NA | 0.30 | 0.30 | <0.2 | <0.2 | 0.3 | 0.4 | 7.4 | 18.9 | 1.6 | 7.5 | 1.9 | 0.8 | 7.5 |
| PCDF3-APA | <0.2 | 160 | NA | 0.40 | <0.20 | <0.2 | <0.2 | 0.4 | 0.3 | 11.9 | 19.9 | 2.5 | 5.1 | 1.4 | 0.8 | 5.1 |
| PFDF4-APA | <0.2 | 100 | NA | 0.70 | 0.70 | <0.2 | <0.2 | <0.2 | 0.3 | 5.5 | 48.6 | 0.2 | 6.4 | 1.2 | 1.0 | 6.4 |
| PLDF1B-APA | <0.2 | 10.0 | NA | 0.40 | 0.90 | <0.2 | <0.2 | 0.3 | 0.3 | 4.6 | 17.0 | 0.8 | 4.6 | 4.3 | 0.8 | 4.6 |
| PLDF2-APA | <0.2 | 10.0 | NA | 0.30 | 1.40 | <0.2 | <0.2 | <0.2 | 0.1 | 2.7 | 19.5 | 0.9 | 4.7 | 4.2 | 0.7 | 4.7 |
| PLDF3-APA | <0.2 | 20.0 | NA | 0.30 | 0.70 | 1.50 | <0.2 | <0.2 | 0.3 | 10.2 | 24.2 | 1.0 | 7.7 | 2.3 | 0.7 | 7.7 |
| PMDF9-APA | <0.2 | 30.0 | NA | 0.20 | 0.50 | 1.00 | <0.2 | <0.2 | 0.7 | 5.1 | 16.9 | 3.3 | 13.5 | 61.5 | 1.9 | 13.5 |

## References

• Adekanmbi, A. A., Shu, X., Zou, Y., and Sizmur, T. (2022). Legacy effect of constant and diurnally oscillating temperatures on soil respiration and microbial community structure. European Journal of Soil Science, 73(6), e13319, 10.1111/ejss.13319

• Akbari, A., and Ghoshal, S. (2015). Effects of diurnal temperature variation on microbial community and petroleum hydrocarbon biodegradation in contaminated soils from a sub-Arctic site. Environmental microbiology, 17(12), 4916–4928, 10.1111/1462-2920.12846

• Amaresan, N., Kumar, K., Venkadesaperumal, G., and Srivathsa, N. C. (2018). Microbial community level physiological profiles of active mud volcano soils in Andaman and Nicobar Islands. National Academy Science Letters, 41, 161–164, 10.25083/rbl/25.4/1731.1736

• Asif, A., Koner, S., Chen, J. S., Hussain, A., Huang, S. W., Hussain, B., & Hsu, B. M. (2024). Uncovering the microbial community structure and physiological profiles of terrestrial mud volcanoes: a comprehensive metagenomic insight towards their Trichloroethylene biodegradation potentiality. Environmental Research, 119457, 10.1016/j.envres.2024.119457

• Barton, H. A., and Northup, D. E. (2007). Geomicrobiology in cave environments: past, current and future perspectives. Journal of Cave and Karst Studies, 69(1), 163–178.

• Barton, H. A., Giarrizzo, J. G., Suarez, P., Robertson, C. E., Broering, M. J., Banks, E. D., and Venkateswaran, K. (2014). Microbial diversity in a Venezuelan orthoquartzite cave is dominated by the Chloroflexi (Class Ktedonobacterales) and Thaumarchaeota Group I. 1c. Frontiers in microbiology, 5, 615, 10.3389/fmicb.2014.00615

• Bleahu, M., Decu, V., Negrea, S., Pleşa, C., Povară, I., and Viehmann, I. (1976). Caves from Romania. Editura Ştiințifică şi Enciclopedică, Bucharest.

• Bogdan D.F., Baricz A.I., Chiciudean I., Bulzu P.A., Cristea A., Năstase-Bucur R., Levei E.A., Cadar O., Sitar C., Banciu H.L. and Moldovan O.T., (2023), Diversity, distribution and organic substrates preferences of microbial communities of a low anthropic activity cave in North-Western Romania. Frontiers in Microbiology 14:962452, 10.3389/fmicb.2023.962452

• Boteva, S., Stanachkova, M., Traykov, I., Angelova, B., & Kenarova, A. (2024). Bacterial functional responses to environmental variability: a case study in three distinct mountain lakes within a single watershed. Biotechnology & Biotechnological Equipment, 38(1), 2418549, 10.1080/13102818.2024.2418549

• Bücs, S. L., Jére, C., Csősz, I., Barti, L., and Szodoray-Parády, F. (2012). Distribution and conservation status of cave-dwelling bats in the Romanian Western Carpathians. Vespertilio 16, 97–113.

• Cacchio, P., and Del Gallo, M. (2019). A novel approach to isolation and screening of calcifying bacteria for biotechnological applications. Geosciences, 9(11), 479, 10.3390/geosciences9110479

• Cristea, A., Andrei, A. Ș., Baricz, A., Muntean, V., Banciu, H. L. (2014): Rapid assessment of carbon substrate utilization in the epilimnion of meromictic Ursu Lake (Sovata, Romania) by the Biolog Eco Plate™ approach. Studia Universitatis Babes-Bolyai, Biologia 59(1): 41–53

• D’Angeli, I. M., Ghezzi, D., Leuko, S., Firrincieli, A., Parise, M., Fiorucci, A., and Cappelletti, M. (2019). Geomicrobiology of a seawater-influenced active sulfuric acid cave. PLoS One, 14(8), e0220706, 10.1371/journal.pone.0220706.

• De Mandal, S., Chatterjee, R., and Kumar, N. S. (2017). Dominant bacterial phyla in caves and their predicted functional roles in C and N cycle. BMC Microbiol=ogy, 17:90. 10.1186/s12866-017-1002-x

• Dencker, T. S., Pecuchet, L., Beukhof, E., Richardson, K., Payne, M. R., and Lindegren, M. (2017). Temporal and spatial differences between taxonomic and trait biodiversity in a large marine ecosystem: Causes and consequences. PLoS One, 12(12), e0189731, 10.1371/journal.pone.0189731

• Epure, L., Meleg, I. N., Munteanu, C. M., Roban, R. D., and Moldovan, O. T. (2014). Bacterial and fungal diversity of quaternary cave sediment deposits. Geomicrobiology Journal 31, 116–127, 10.1080/01490451.2013.815292

• Feigl, V., Ujaczki, É., Vaszita, E., and Molnár, M. (2017). Influence of red mud on soil microbial communities: Application and comprehensive evaluation of the Biolog^®^ EcoPlate™ approach as a tool in soil microbiological studies. Science of the Total Environment, 595, 903–911, 10.1016/j.scitotenv.2017.03.266

• Garland, J. L. (1997). Analysis and interpretation of community-level physiological profiles in microbial ecology. FEMS Microbiology ecology, 24(4), 289–300, 10.1111/j.1574-6941.1997.tb00446.x

• Ghaly, T. M., Focardi, A., Elbourne, L. D. H., et al. (2023). Stratified microbial communities in Australia’s only anchialine cave are taxonomically novel and drive chemotrophic energy production via coupled nitrogen-sulphur cycling. Microbiome, 11, 190. 10.1186/s40168-023-01633-8

• Gryta, A., Frąc, M., and Oszust, K. (2014). The application of the Biolog EcoPlate approach in ecotoxicological evaluation of dairy sewage sludge. Applied biochemistry and biotechnology, 174, 1434–1443, 10.1007/s12010-014-1131-8

• Haegeman, B., Hamelin, J., Moriarty, J., Neal, P., Dushoff, J., and Weitz, J. S. (2013). Robust estimation of microbial diversity in theory and in practice. The ISME journal, 7(6), 1092–1101, 10.1038/ismej.2013.10

• Inskeep, W. P., Jay, Z. J., Tringe, S. G., Herrgård, M. J., Rusch, D. B., and YNP Metagenome Project Steering Committee and Working Group Members. (2013). The YNP metagenome project: environmental parameters responsible for microbial distribution in the Yellowstone geothermal ecosystem. Frontiers in microbiology, 4, 67, 10.3389/fmicb.2013.00067

• Iţcuş, C., Pascu, M. D., Brad, T., Perşoiu, A., and Purcarea, C. (2016). Diversity of cultured bacteria from the perennial ice block of Scarisoara Ice Cave, Romania. International journal of speleology, 45(1), 9, 10.5038/1827-806X.45.1.1948

• Jałowiecki, Ł., Chojniak, J. M., Dorgeloh, E., Hegedusova, B., Ejhed, H., Magnér, J., and Płaza, G. A. (2016). Microbial community profiles in wastewaters from onsite wastewater treatment systems technology. PloS one, 11(1), e0147725, 10.1371/journal.pone.0147725

• Jiang, Y., Wang, Y., Huang, Z., Zheng, B., Wen, Y., and Liu, G. (2023). Investigation of phytoplankton community structure and formation mechanism: a case study of Lake Longhu in Jinjiang. Frontiers in Microbiology, 14, 1267299, 10.3389/fmicb.2023.1267299.

• Jin, Q., Black, A., Kales, S. N., Vattem, D., Ruiz-Canela, M., and Sotos-Prieto, M. (2019). Metabolomics and Microbiomes as Potential Tools to Evaluate the Effects of the Mediterranean Diet. Nutrients, 11(1), 207, 10.3390/nu11010207

• Jones, D. (2015). 2. Methods for Characterizing Microbial Communities in Caves and Karst: A Review. In A. Summers Engel (Ed.), Microbial Life of Cave Systems (pp. 23-46). Berlin, München, Boston: De Gruyter, 10.1515/9783110339888-004

• Koner, S., Chen, J. S., Hsu, B. M., Rathod, J., Huang, S. W., Chien, H. Y., … and Chan, M. W. (2022). Depth-resolved microbial diversity and functional profiles of trichloroethylene-contaminated soils for Biolog EcoPlate-based biostimulation strategy. Journal of hazardous materials, 424, 127266, 10.1016/j.jhazmat.2021.127266

• Koner, S., Chen, J. S., Hsu, B. M., Tan, C. W., Fan, C. W., Chen, T. H., and Nagarajan, V. (2021). Assessment of Carbon substrate catabolism pattern and functional metabolic pathway for Microbiota of Limestone Caves. Microorganisms, 9, 10.3390/microorganisms9081789.

• Li, H., Xu, Z., Yan, Q., Yang, S., Van Nostrand, J. D., Wang, Z., and Deng, Y. (2018). Soil microbial beta-diversity is linked with compositional variation in aboveground plant biomass in a semi-arid grassland. Plant and Soil, 423, 465–480, 10.1007/s11104-017-3524-2.

• Li, X., Liu, L., Zhu, Y., Zhu, T., Wu, X., and Yang, D. (2021). Microbial community structure and its driving environmental factors in black carp (Mylopharyngodon piceus) aquaculture pond. Water, 13(21), 3089, 10.3390/w13213089

• Liu, F., Giometto, A., and Wu, M. (2021). Microfluidic and mathematical modeling of aquatic microbial communities. Analytical and Bioanalytical Chemistry, 413, 2331– 2344, 10.1007/s00216-020-03085-7

• Liu, L., Li, A., Zhu, L., Xue, S., Li, J., Zhang, C., and Mao, Y. (2023). The Application of the Generalized Additive Model to Represent Macrobenthos near Xiaoqing Estuary, Laizhou Bay. Biology, 12(8), 1146, 10.3390/biology12081146

• Martin-Pozas, T., Gonzalez-Pimentel, J. L., Jurado, V., Cuezva, S., Dominguez-Moñino, I., Fernandez-Cortes, A., and Saiz-Jimenez, C. (2020). Microbial activity in subterranean ecosystems: Recent advances. Applied Sciences, 10(22), 8130, 10.3390/app10228130

• Melita, M., Amalfitano, S., Preziosi, E., Ghergo, S., Frollini, E., Parrone, D., and Zoppini, A. (2023). Redox conditions and a moderate anthropogenic impairment of groundwater quality reflected on the microbial functional traits in a volcanic aquifer. Aquatic Sciences, 85(1), 3, 10.1007/s00027-022-00899-8

• Meyer, N. R., Parada, A. E., Kapili, B. J., Fortney, J. L., and Dekas, A. E. (2022). Rates and physico-chemical drivers of microbial anabolic activity in deep-sea sediments and implications for deep time. Environmental Microbiology, 24(11), 5188–5201, 10.1111/1462-2920.16183

• Moldovan, O. T., Carrell, A. A., Bulzu, P. A., Levei, E., Bucur, R., Sitar, C., and Podar, M. (2023). The gut microbiome mediates adaptation to scarce food in Coleoptera. Environmental Microbiome, 18(1), 80, 10.1186/s40793-023-00537-2

• Moretti, G., Matteucci, F., Ercole, C., Vegliò, F., and Del Gallo, M. (2016). Microbial community distribution and genetic analysis in a sludge active treatment for a complex industrial wastewater: A study using microbiological and molecular analysis and principal component analysis. Annals of microbiology, 66, 397–405, 10.1007/s13213-015-1122-1

• Morita, N., Toma, Y., and Ueno, H. (2024). Microbial diversity and community structure in co-composted bamboo powder and tea leaves based on carbon substrate utilization patterns of the BIOLOG EcoPlate method. Advances in Bamboo Science, 8, 100101, 10.1016/j.bamboo.2024.100101

• Musat, N., Musat, F., Weber, P. K., and Pett-Ridge, J. (2016). Tracking microbial interactions with NanoSIMS. Current Opinion in Biotechnology, 41, 114–121, 10.1016/j.copbio.2016.06.007

• Nyyssönen, M., Hultman, J., Ahonen, L., Kukkonen, I., Paulin, L., Laine, P., … and Auvinen, P. (2014). Taxonomically and functionally diverse microbial communities in deep crystalline rocks of the Fennoscandian shield. The ISME journal, 8(1), 126–138, 10.1038/ismej.2013.125

• Obusan, M. C. M., Castro, A. E., Villanueva, R. M. D., Isagan, M. D. E., Caras, J. A. A., and Simbahan, J. F. (2023). Physico-Chemical Quality and Physiological Profiles of Microbial Communities in Freshwater Systems of Mega Manila, Philippines. Data, 8(6), 103, 10.3390/data8060103

• O’Connor, B. R., Fernández-Martínez, M. Á., Léveillé, R. J., & Whyte, L. G. (2021). Taxonomic characterization and microbial activity determination of cold-adapted microbial communities in Lava Tube Ice Caves from Lava Beds National Monument, a high-fidelity Mars analogue environment. Astrobiology, 21(5), 613–627, 10.1089/ast.2020.2327

• Park, S., Cho, Y. J., Jung, D. Y., Jo, K. N., Lee, E. J., and Lee, J. S. (2020). Microbial diversity in moonmilk of Baeg-nyong Cave, Korean CZO. Frontiers in Microbiology, 11, 613, 10.3389/fmicb.2020.00613

• Pašić, L., Kovče, B., Sket, B., and Herzog-Velikonja, B. (2009). Diversity of microbial communities colonizing the walls of a Karstic cave in Slovenia. FEMS Microbiology Ecology, 71(1), 50–60, 10.1111/j.1574-6941.2009.00789.x

• Patsch, D., van Vliet, S., Marcantini, L. G., and Johnson, D. R. (2018). Generality of associations between biological richness and the rates of metabolic processes across microbial communities. Environmental microbiology, 20(12), 4356–4368, 10.1111/1462-2920.14352

• Paula, C. C. P. D., Bichuette, M. E., and Seleghim, M. H. R. (2020). Nutrient availability in tropical caves influences the dynamics of microbial biomass. MicrobiologyOpen, 9(7), e1044, 10.1002/mbo3.1044

• Pedersen, K. (2012). Subterranean microbial populations metabolize hydrogen and acetate under in situ conditions in granitic groundwater at 450 m depth in the Äspö Hard Rock Laboratory, Sweden. FEMS Microbiology Ecology, 81(1), 217–229. 10.1111/j.1574-6941.2012.01370.x

• Power, J. F., Carere, C. R., Lee, C. K., Wakerley, G. L., Evans, D. W., Button, M., … and Stott, M. B. (2018). Microbial biogeography of 925 geothermal springs in New Zealand. Nature communications, 9(1), 2876, 10.1038/s41467-018-05020-y

• Rutgers, M., Wouterse, M., Drost, S. M., Breure, A. M., Mulder, C., Stone, D., … & Bloem, J. (2016). Monitoring soil bacteria with community-level physiological profiles using Biolog™ ECO-plates in the Netherlands and Europe. Applied Soil Ecology, 97, 23–35, 10.1016/j.apsoil.2015.06.007

• Shaw, A. K., Halpern, A. L., Beeson, K., Tran, B., Venter, J. C., and Martiny, J. B. (2008). It’s all relative: ranking the diversity of aquatic bacterial communities. Environmental microbiology, 10(9), 2200–2210, 10.1111/j.1462-2920.2008.01626.x

• Shen, J., Smith, A. C., Barnett, M. J., Morgan, A., and Wynn, P. M. (2022). Distinct microbial communities in the soils, waters, and speleothems of a hyperalkaline cave system. Journal of Geophysical Research: Biogeosciences, 127(9), e2022JG006866

• Shen, J., Smith, A. C., Barnett, M. J., Morgan, A., and Wynn, P. M. (2022). Distinct microbial communities in the soils, waters, and speleothems of a hyperalkaline cave system. Journal of Geophysical Research: Biogeosciences, 127(9), e2022JG006866, 10.3390/microorganisms9081789

• Simpson, G. L., (2024). gratia: An R package for exploring generalized additive models. Journal of Open Source Software, 9(104), 6962, 10.21105/joss.06962

• Sofo, A., and Ricciuti, P. (2019). A standardized method for estimating the functional diversity of soil bacterial community by Biolog® EcoPlatesTM assay—the case study of a sustainable olive orchard. Applied Sciences, 9(19), 4035, 10.3390/app9194035

• Stefanowicz, A. (2006). The Biolog plates technique as a tool in ecological studies of microbial communities. Polish Journal of Environmental Studies, 15(5).

• Tang, Z., Sun, X., Luo, Z., He, N., and Sun, O. J. (2018). Effects of temperature, soil substrate, and microbial community on carbon mineralization across three climatically contrasting forest sites. Ecology and Evolution, 8(2), 879–891, 10.1002/ece3.3708

• Teng, Z., Fan, W., Wang, H., Cao, X., & Xu, X. (2020). Monitoring soil microorganisms with community-level physiological profiles using Biolog EcoPlates™ in Chaohu lakeside wetland, East China. Eurasian Soil Science, 53, 1142–1153, 10.1134/S1064229320080141

• Theodorescu, M., Bucur, R., Bulzu, PA., Faur L., Levei EA., Mirea IC, Cadar O., Ferreira R, Souza-Silva M., and Moldovan OT. (2023). Environmental Drivers of the Moonmilk Microbiome Diversity in Some Temperate and Tropical Caves. Microbial Ecology, 86,2847–2857, 10.1007/s00248-023-02286-8

• Thompson, H. F., and Gutierrez, T. (2021). Detection of hydrocarbon-degrading bacteria on deepwater corals of the northeast Atlantic using CARD-FISH. Journal of Microbiological Methods, 187, 106277, 10.1016/j.mimet.2021.106277

• Tobias-Hünefeldt, S. P., Wing, S. R., Baltar, F., and Morales, S. E. (2021). Changes in microbial community phylogeny and metabolic activity along the water column uncouple at near sediment aphotic layers in fjords. Scientific Reports, 11(1), 19303, 10.1038/s41598-021-98519-2

• Wu, X., Holmfeldt, K., Hubalek, V., Lundin, D., Åström, M., Bertilsson, S., and Dopson, M. (2016). Microbial metagenomes from three aquifers in the Fennoscandian shield terrestrial deep biosphere reveal metabolic partitioning among populations. The ISME Journal, 10(5), 1192–1203, 10.1038/ismej.2015.185

• Yu, K., Yi, S., Li, B., et al. (2019). An integrated meta-omics approach reveals substrates involved in synergistic interactions in a bisphenol A (BPA)-degrading microbial community. Microbiome, 7(1), 16. 10.1186/s40168-019-0634-5

• Yu, Z., Yang, J., Amalfitano, S., Yu, X., and Liu, L. (2014). Effects of water stratification and mixing on microbial community structure in a subtropical deep reservoir. Scientific reports, 4(1), 5821, 10.1038/srep05821

• Yun, Y., Cheng, X., Wang, W., and Wang, H. (2018). Seasonal variation of bacterial community and their functional diversity in drip water from a karst cave. Chinese Science Bulletin, 63(36), 3932–3944, 10.1360/N972018-00627

• Zak, J. C., Willig, M. R., Moorhead, D. L., and Wildman, H. G. (1994). Functional diversity of microbial communities: a quantitative approach. Soil Biology and Biochemistry, 26(9), 1101–1108, 10.1016/0038-0717(94)90131-7

• Zhang, P., Cui, Z., Guo, M., and Xi, R. (2020). Characteristics of the soil microbial community in the forestland of Camellia oleifera. Peer J 8: e9117, doi 10.7717/peerj.9117

• Zheng, Y., and Gong, X. (2019). Niche differentiation rather than biogeography shapes the diversity and composition of microbiome of *Cycas panzhihuaensis*. Microbiome, 7, 1–19, 10.1186/s40168-019-0770-y

• Zoltan, L., and Szántó, L. (2003). Bats of the Carpathian Region. Acta Chiropt. 5, 155–160. 10.3161/001.005.0115

